# High-Throughput Identification of Calcium Regulated Proteins Across Diverse Proteomes

**DOI:** 10.1101/2024.01.18.575273

**Authors:** Timothy M. Locke, Rose Fields, Hayden Gizinski, George M. Otto, David M. Shechner, Matthew D. Berg, Judit Villen, Yasemin Sancak, Devin Schweppe

## Abstract

Calcium ions play important roles in nearly every biological process, yet whole-proteome analysis of calcium effectors has been hindered by lack of high-throughput, unbiased, and quantitative methods to identify proteins-calcium engagement. To address this, we adapted protein thermostability assays in the budding yeast, human cells, and mouse mitochondria. Based on calcium-dependent thermostability, we identified 2884 putative calcium-regulated proteins across human, mouse, and yeast proteomes. These data revealed calcium engagement of novel signaling hubs and cellular processes, including metabolic enzymes and the spliceosome. Cross-species comparison of calcium-protein engagement and mutagenesis experiments identified residue-specific cation engagement, even within well-known EF-hand domains. Additionally, we found that the dienoyl-CoA reductase DECR1 binds calcium at physiologically-relevant concentrations with substrate-specific affinity, suggesting direct calcium regulation of mitochondrial fatty acid oxidation. These unbiased, proteomic analyses of calcium effectors establish a key resource to dissect cation engagement and its mechanistic effects across multiple species and diverse biological processes.

## INTRODUCTION

Calcium ions (Ca^2+^) are essential for life. They play important signaling, catalytic and structural roles through their interactions with proteins. These Ca^2+^-binding proteins regulate diverse biological processes across species. In yeast, Ca^2+^-signaling mediates transcription^1^ and cell death^2,3^. In mammals, Ca^2+^ regulates a wide range of events, from fertilization to insulin secretion to muscle contraction. In addition, organelle-specific calcium-signaling has been long appreciated to coordinate organellar functions. For example, mitochondrial calcium-binding proteins play established roles in activating the TCA cycle, regulating metabolite and ion carriers, and opening the permeability transition pore^4,5^. Mitochondrial fatty acid oxidation, amino acid metabolism, protein homeostasis, and electron transport chain activity^6–9^ are also Ca^2+^-regulated. A larger scope of Ca^2+^ affected biological processes in multiple species has been suggested for a plethora of phenotypes^10–13^–often without clear molecular mechanisms–suggesting the presence of as yet unidentified Ca^2+^-regulated proteins. Taken together, the lack of a comprehensive atlas of Ca^2+^-binding proteins across common model organisms creates a major challenge when investigating molecular mechanisms underlying Ca^2+^-regulated biology.

Currently, there are 720 proteins annotated as Ca^2+^ ion binding in the human proteome (GO:0005509; 3.53% of 20,385 human UniProt entries). Traditionally, Ca^2+^-binding proteins have been identified through computational prediction of well-defined binding motifs (e.g., EF-hands) or low-throughput biochemical analysis of highly purified proteins using specialized methods such as isothermal calorimetry. However, the majority of Ca^2+^-binding proteins do not have a defined ion-binding motif: only 10% of known Ca^2+^-binding sites in the Protein Data Bank (PDB) contain EF-Hand-like domains^14^. Instead, most calcium-binding proteins coordinate the ion by amino acid functional groups that come together in the folded protein to form a binding pocket. Computational efforts to predict potential Ca^2+^ and other metal binding pockets have been developed^15^, but their widespread application has been hindered by their lack of specificity for metal binding and poor precision^16^. Structure prediction using AlphaFold and computational ligand binding prediction with AlphaFill^17^ provide binding predictions for a range of ligands including Ca^2+^, but the specificity and accuracy of this approach has not been tested for Ca^2+^-binding. As a result, despite governing all aspects of biology, Ca^2+^-binding proteins remain hard to identify using available methods.

Thermal proteome profiling has emerged as a powerful assay to detect protein engagement of ligands and small-molecules^18^ by leveraging the fact that each protein has a characteristic melting temperature that is altered upon ligand binding. In this assay, proteins in cells or lysates are exposed to a temperature gradient. Proteins that remain soluble at each temperature are quantified and compared, enabling generation of melting curves for thousands of proteins within a single experiment. These assays were initially applied to determine protein engagement of small-molecule drugs^18^, and have since have been applied to identify protein-protein interactions^19^ and nonspecific ligands such as lactate^20^ and ATP^21^. Full melting curve thermal stability assays have previously been used to identify protein-Ca^2+^ engagement at concentrations up to 1mM in the prokaryotic parasite *Toxoplasma gondii*^22^. However, these canonical assays require quantitative measurement of a protein at each individual temperature point to generate accurate melting curves, reducing sample throughput^18^. Thermal proteome profiling assay throughput can be increased by pooling of the temperature points to ‘integrate’ the soluble protein abundance across temperatures into a single point measurement using a method termed protein integral solubility alteration (PISA) assay^23^. PISA enables collection, quantification, and comparison of multiple replicates of each ligand condition to improve throughput and enable the usage of common methods for statistical comparison of protein populations and the analysis of differential protein abundance^23–25^.

Here, we adapted the PISA assay^23,25^ for unbiased and high-throughput identification of Ca^2+^-regulated proteins (CRPs) from diverse biological samples. We reasoned that PISA could be used to efficiently identify CRPs across multiple species and conditions. PISA could thereby help to overcome current limitations for the identification of Ca^2+^-binding proteins and uncover proteins that are indirectly regulated by Ca^2+^ via secondary interactions with direct Ca^2+^ binders. Using EGTA as a Ca^2+^-specific chelator and Mg^2+^ as a control for cation addition, we coupled PISA with isobaric sample multiplexing to quantify Ca^2+^ engagement across proteomes. Our multispecies datasets generated proteome-scale insights of Ca^2+^ binding and signaling. Proteins exhibiting a Ca^2+^-dependent thermal stability shift in our datasets were enriched for known Ca^2+^-binding proteins, validating the ability of our approach to identify novel Ca^2+^ interactors. Cross-species analysis allowed us to identify specific amino acid substitutions in highly conserved proteins that confer ion specificity. Additionally, we identified DECR1, an enzyme involved in the oxidation of polyunsaturated fatty acids (PUFA), as a novel Ca^2+^-binding mitochondrial protein^26,27^. Together this work establishes the utility of thermal stability proteomics for the biochemical identification of CRPs. We note that these methods could be readily deployed in the future to identify protein engagement for diverse groups of ligands, including metal cations, co-factors, and enzymatic substrates.

## RESULTS

### Optimization of a thermal stability workflow to identify Ca^2+^-regulated proteins

Identifying ligand-protein interactions for non-specific ligands can be challenging^20,21^. To induce measurable quantitative changes in stability generally requires a clean comparative background and excess ligand to induce measurable effect sizes even when individual affinities are low^20^. We quantified Ca^2+^-engagement in proteomes of three diverse model systems: yeast, human, and mice. Owing to the large relative disparity in the number of annotated Ca^2+^-binding proteins in UniProt in yeast (71 of 6,727 proteins, 1%) and human (1,232 of 20,428 proteins, 6%) cells, we measured Ca^2+^-dependent thermal stability in crude extracts from human HEK293T cells and *Saccharomyces cerevisiae* (BY4742) cells. We additionally sought to determine if we could measure Ca^2+^-binding within the context of organelles. Based on extensive phenotypic data relating Ca^2+^ to mitochondrial processes^28,29^, we investigated Ca^2+^-dependent thermal stability alteration *in situ* in murine mitochondria.

Native proteomes were isolated from yeast and human cells and murine mitochondria were enriched from livers. All samples were prepared in a buffer that contained 5mM EGTA, a high affinity Ca^2+^ chelator, to significantly decrease the concentration of free Ca^2+^ and strip metal-binding proteins in the untreated lysate (**Fig. 1A**). The whole-proteome human and yeast studies were performed in cell lysates as previously described^25^. The mitochondrial specific mouse study was adapted from Schweppe et al., 2017 and Frezza et al., 2007^30,31^. The Ca^2+^-reduced lysate was then divided into four samples: the first receiving no additional treatment, and the remaining three receiving addition of either 10 mM CaCl_2_, 5 mM MgCl_2_ or both 10mM CaCl_2_ and 5mM MgCl_2_. The MgCl_2_ condition was added to control for non-specific divalent cation binding and owing to the specificity of EGTA for Ca^2+^, less MgCl_2_ was required to achieve near physiological concentrations of Mg^2+^ (**Fig. S1A**). Additionally, we conjectured that treatment with both Ca^2+^ and Mg^2+^ might uncover Mg^2+^-dependent Ca^2+^-binding events in our samples^32^. Our sequential chelation and ion addition strategy enabled better control of free ion concentration in the experimental samples. After EGTA addition, we measured the free Ca^2+^ and Mg^2+^ concentrations to be lower than their free concentrations in resting cells (∼100 nM for Ca^2+^and ∼1 mM for Mg^2+^), allowing the formation of the unbound form of Ca^2+^-or Mg^2+^-binding proteins^33^ (**Fig. S1A**). After cation addition, the free concentrations increased to levels that should saturate cation binding domains and favor cation-protein interactions (**Fig. S1A**)^34^. Under these conditions, we quantified the relative abundance of proteins following the experimental flow shown in **Fig 1A**. To identify Calcium Regulated Proteins (CRPs), we established a definition for significant thermal stability changes and hit criteria (**Fig. S1B**). A protein was determined to be a CRP if it had a significant thermal stability change in response to Ca^2+^ and passing through two filtering steps: lack of Mg^2+^ engagement (Class 1), or significant Ca^2+^engagement in the presence of Mg^2+^ (Class 2). Through this criteria, non-specific cation responsive proteins were separated from identified CRPs.

**Figure 1:**
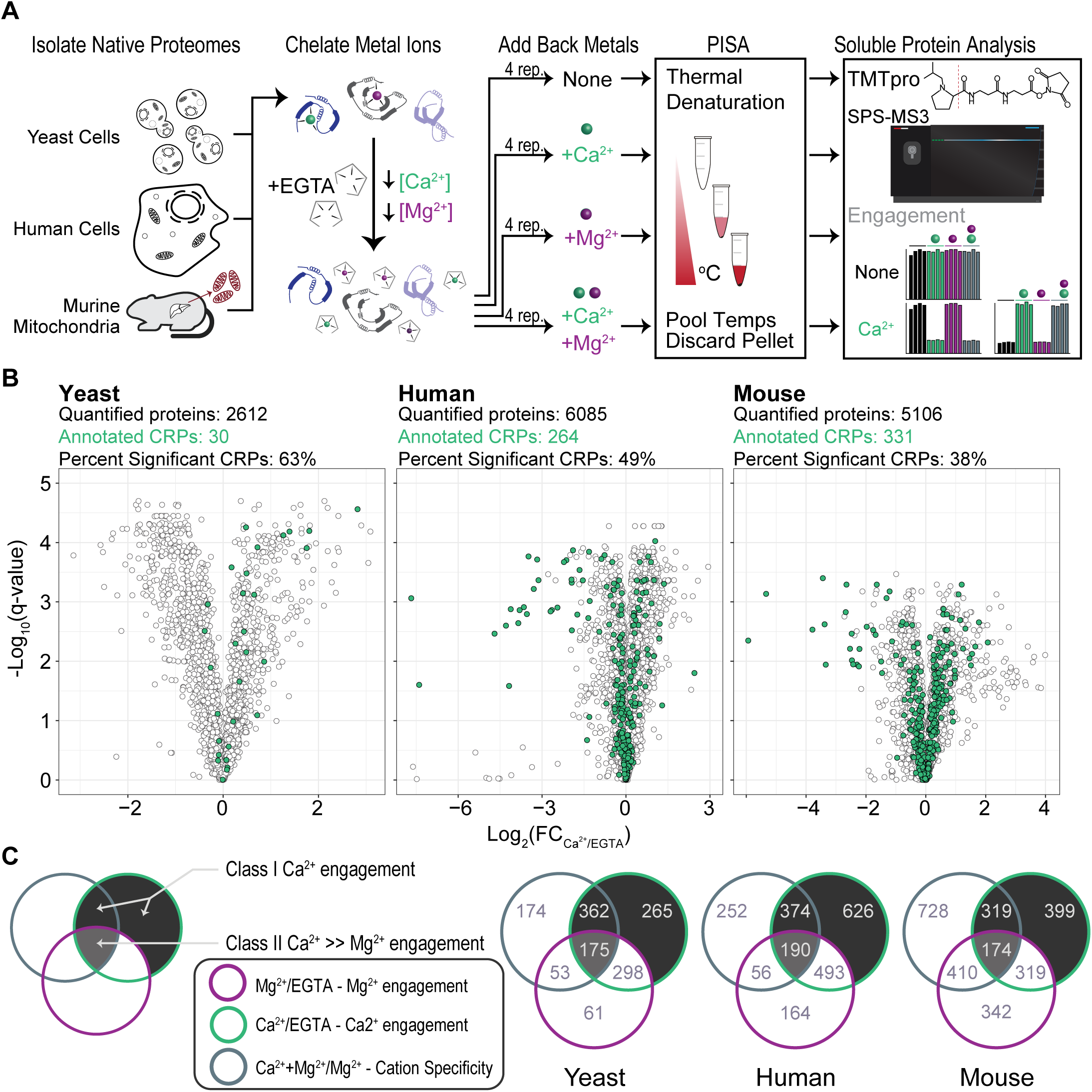
PISA with isobaric sample multiplexing in three different proteomes identifies Ca^2+^-regulated proteins. **(A)** Schematics of the experimental workflow to detect ion-induced changes in protein thermal stability. Lysates from yeast, human and isolated murine liver mitochondria were treated with chelator EGTA. After the addition of ions, a temperature gradient was applied to aliquoted samples, the samples were then mixed, and insoluble (heat denatured) material was precipitated. Soluble proteins were labeled with isobaric tags, and protein abundance was quantified. **(B)** Volcano plots show protein abundance in Ca^2+^ relative to EGTA, and their q-values. Proteins with a |log2(FC)| ≥ 0.2, and q-value ≤ 0.05 (Welch’s t-test with multiple hypothesis correction) were considered to show significant thermal stability changes. Green dots indicate proteins with previous Ca^2+^ annotation. **(C)** Venn diagrams show proteins with significant thermal stability changes in each category listed. Proteins that are in the shaded compartments represent CRPs. Proteins that do not show a significant thermal stability change between Mg^2+^ and Mg^2+^Ca^2+^ conditions were considered to be non-specific ion responders and excluded from the list of CRPs.

### Ca^2+^-dependent thermal stability across the proteomes enriches known Ca^2+^-regulated pathways

From the three thermal stability analyses, we quantified 2,612 yeast proteins, 6,085 human proteins, and 5,106 mouse proteins (**Fig. 1B, Fig. S1C**). Principal component analysis resulted in clear differentiation of conditional effects based on cation treatment (**Fig. S1D**). When we applied our hit criteria, we observed Ca^2+^-specific changes in thermal stability for 30 known yeast CRPs, 264 known human CRPs and 331 known mouse CRPs. In addition, we identified a total of 802 yeast, 1190 human, and 892 mouse putative CRPs (**Fig. 1B, 1C, Table 1**). The majority of these CRPs had Ca^2+^-dependent thermal stability shifts with no corresponding change in stability due to Mg^2+^ (**Fig. 1C**; Class I CRPs, see Methods). Another subset of proteins had both significant Mg^2+^-dependent and Ca^2+^-dependent thermal stability changes. From the co-treatment comparative samples, we observed proteins with significant thermal stability in the dual cation condition compared to Mg^2+^ suggesting Ca^2+^-specific engagement (Class II CRPs, see Methods). For this work, we aggregated the Class I and Class II CRPs together to perform all subsequent analyses.

**Table 1.** Summary statistics of quantified proteins called CRPs in each of the three thermal stability datasets.

| Sample | Proteins in Proteome | Annotated CRPs in the Proteome | Proteins Quantified | Annotated CRPs Quantified | Annotated CRPs Identified as Hits | Total Called CRPs | Class I CRPs | Class II CRPs |
| --- | --- | --- | --- | --- | --- | --- | --- | --- |
| **Yeast** | 6727 | 71 | 2612 | 30 | 19 (**63.3%**) | 802 (30.5%) | 627 | 175 |
| **Human** | 20428 | 1232 | 6085 | 264 | 129 (**48.9%**) | 1190 (19.6%) | 1000 | 190 |
| **Mouse** | 17184 | 1088 | 5106 | 331 | 127 (**38.3%**) | 892 (17.5%) | 718 | 174 |

We analyzed enriched gene sets in our human and mouse CRPs based on Gene Ontology (GO) Molecular Function (MF) terms (q-value ≤ 0.05; **Figs. 2A, 2B**)^35^. These sets were enriched for terms such as Ca^2+^ ion binding (GO:0005516) and Ca^2+^-dependent phospholipid binding (GO:0005544). We further analyzed proteins with a significant thermal stability change (absolute log_2_ fold change ≥ 0.2 and q-value ≤ 0.05) upon Mg^2+^ addition. The Mg^2+^-protein set returned no Ca^2+^-related terms (**Fig. 2C**) and instead enriched for terms related to DNA binding, RNA binding, and nucleotide coordination, consistent with the role of Mg^2+^ in the binding and stabilization of nucleic acids^36^. The ion-specific enrichments in our hits reinforced the utility of thermal stability assays in differentiating the effects of common divalent cations on complex proteomes.

**Figure 2:**
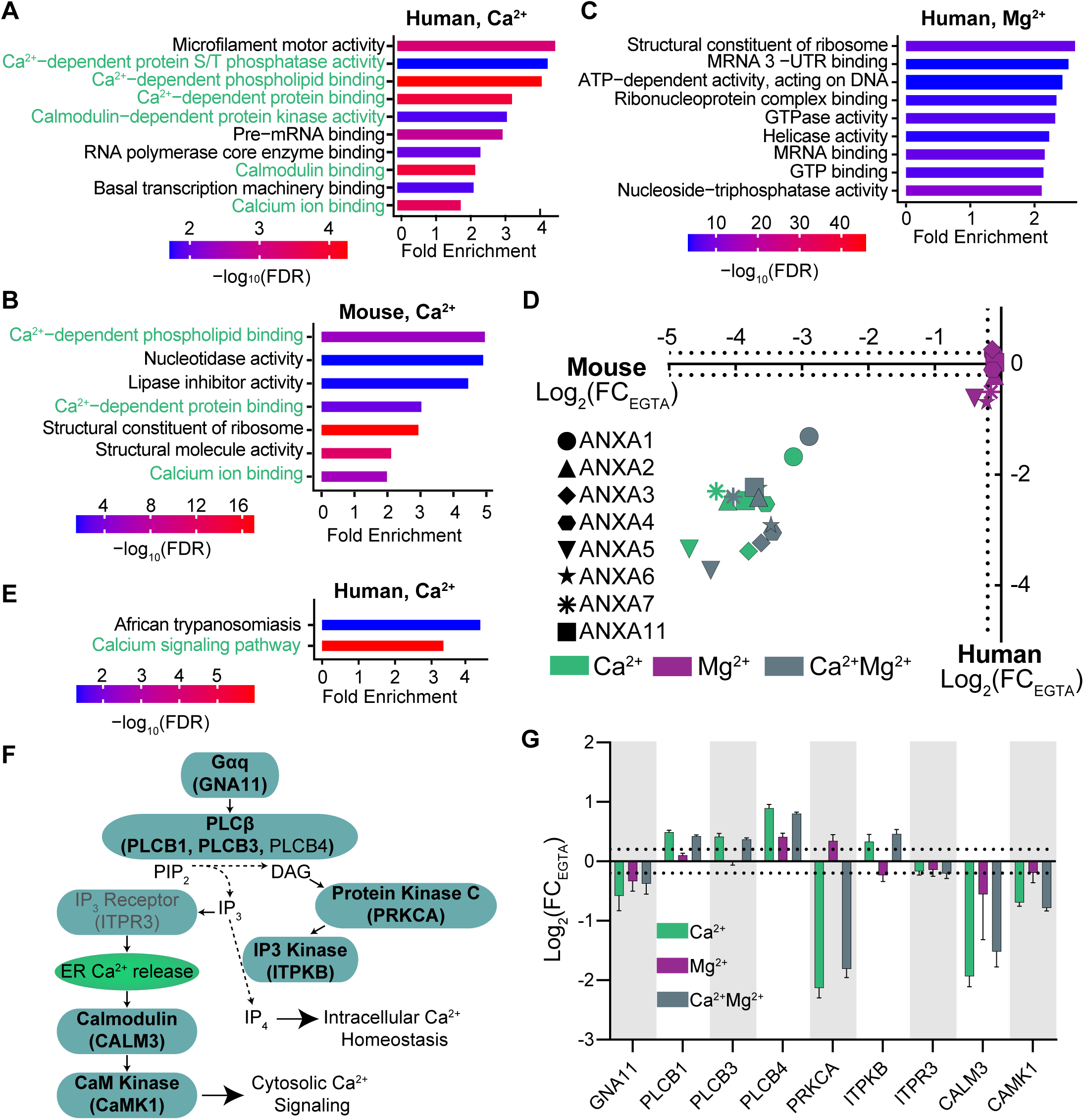
Analysis of human and mouse datasets show significant enrichment of known Ca^2+^-regulated processes, proteins and pathways. **(A,B)** Gene set enrichment analysis (GSEA) of CRPs from human **(A)** and mouse **(B)** datasets show significant enrichment of molecular function terms related to Ca^2+^ binding and signaling (green text). **(C)** GSEA of proteins with a significant thermal stability change in response to Mg^2+^ addition shows an enrichment for DNA, RNA and nucleotide binding molecular functions, and lack Ca^2+^-related processes. **(D)** Ion-induced thermal stability changes of Annexin proteins from human and mouse datasets show a Ca^2+^-specific response. **(E)** GSEAof human dataset using KEGG annotations enriches Ca^2+^ signaling. **(F)** Schematic representation of the Ca^2+^ regulated Gq signaling pathway. Proteins shown were quantified in the human lysates. Bolded proteins are identified as CRPs. **(G)** Ion-induced thermal stability changes of Gq pathway proteins, relative to EGTA condition. Dotted lines show log_2_(FC) ± 0.2. Error bars: standard deviation of replicate measurements.

We next focused on known CRPs, pathways, and protein interactions. Annexins are Ca^2+^-dependent phospholipid binding proteins^37^. We quantified 8 of 12 and 9 of 12 human and mouse annexin proteins, respectively. All of these proteins had significant destabilization in Ca^2+^ and Ca^2+^Mg^2+^ conditions (**Fig. 2D**). In addition, Gene Set Enrichment Analysis (GSEA) of human CRPs using Kyoto Encyclopedia of Genes and Genomes (KEGG) biological pathway annotations were significantly enriched for “calcium signaling pathway (hsa04020)” (3.3 fold enrichment, -log_10_(FDR)=5.96) (**Fig. 2E**). We quantified 39 of the 253 proteins in this calcium signaling pathway and determined that 24 of the 39 were CRPs. Within the core G-protein coupled receptor q (Gq) signaling cascade, Ca^2+^ is used as a second messenger (**Fig. 2F**). We identified 7 of the 9 key components of this pathway as CRPs in human lysates (**Fig. 2G**). Of note, in addition to the direct Ca^2+^-binding protein calmodulin (CaM), we identified its binding partner Ca^2+^/calmodulin-dependent protein kinase 1 (CaMK1) as a CRP. Our data captures Ca^2+^-dependent protein interactions like CaM-CaMK1, highlighting a unique strength of our method in identifying components of ion-regulated protein complexes and signaling cascades. Overall, these data suggest that the Ca^2+^-dependent thermal protein stability assay is a powerful tool for the unbiased identification of CRPs and pathways.

### Ca^2+^-dependent thermal stability analysis of murine mitochondria identifies regulators of mitochondrial Ca^2+^ homeostasis

To test if our CRP thermal stability assays could be extended to in situ analyses of protein-metal engagement, we examined thermal stability in mitochondria enriched from murine livers^30^. As noted above, we focused on the mitochondria due to the well-established roles of mitochondrial Ca^2+^ in cellular energy metabolism and cell death^38^ and additional functions attributed to mitochondrial Ca^2+^ signaling. We used crude preparations to ensure that we maintained mitochondrial structure and integrity during sample processing.

In total we quantified 80% of the MitoCarta mouse mitochondrial proteome (909 of 1,140) and 93.6 % of the annotated liver mitochondrial proteins (650 of 694)^39,40^, consistent with a crude isolation of these organelles^30^ (**Table S3**). Of the 5106 proteins quantified in crude mitochondrial preparations, we observed significant, Ca^2+^-dependent thermal stability shifts for 892 proteins. Similar to the human lysate data, we observed enrichment for Ca^2+^-related molecular functions (**Fig. 2B**)^41,42^. Murine mitochondria contains 50 known annotated CRPs, with 16 EF-hand-containing proteins^43^, three TCA cycle components^4,5^, and HSP90 ^8^, as well as proteins that associate with Ca^2+^-binding proteins without directly binding Ca^2+^ themselves, such as EMRE^44^. We quantified 23/50 of these proteins in murine mitochondria and called 11 of them as CRPs (47%), compared with 17.5% of all proteins in this dataset. These CRPs included LETM1 and members of the mitochondrial Ca^2+^ uniporter function (MICU1, MICU2), proteins that regulate mitochondrial Ca^2+^ homeostasis^38,45^. In the human dataset, we identified 21 known mitochondrial CRPs and found 7 as CRPs (33%) **(Tables S2)**. Strongly, we observed 101 human mitochondrial CRPs and 236 mouse mitochondrial CRPs suggesting a larger role for mitochondrial Ca^2+^ as a signaling molecule in mammalian systems than previously investigated.

### The spliceosome is enriched with Ca^2+^-regulated proteins

To identify novel Ca^2+^-regulated biological processes, we focused on human CRPs that did not have a previous Ca^2+^-annotation. GSEA of these CRPs based on the KEGG database revealed an unexpected enrichment of proteins that function in the spliceosome (hsa03040) (**Fig. S2A**). The spliceosome is composed of five small nuclear ribonucleoprotein (snRNP) complexes (U1, U2, U4, U5,U6) and numerous additional non-snRNP proteins that all together orchestrate pre-mRNA splicing^46^. We identified 118 of the 129 spliceosomal proteins in our human dataset, and 36 of these have Ca^2+^-dependent thermal stability shifts (**Fig. S2B**). Of the top 10 proteins with the largest thermal stability changes with Ca^2+^ (**Figure S2C**), RBM22 has been shown to be regulated by the EF-hand containing protein ALG2^47^. CHERP, another Ca^2+^-dependent ALG2 interacting protein is also one of the spliceosome CRPs (**Fig. S3B**)^48^. Regulation of alternative splicing by Ca^2+^ signaling through CaMK IV and the CaMK IV-responsive RNA element (CaRRE)^49,50^ has been shown. Our data are consistent with previous literature and suggest that Ca^2+^ ions play a previously underappreciated structural, catalytic, or regulatory role in the function of the core splicing machinery.

### Cross-species whole proteome quantification of Ca^2+^ engagement in human and yeast proteomes identifies conserved and distinct Ca^2+^-binding sites across evolutionary time

Quantification of divalent cation thermal stability experiments across multiple species provided a unique opportunity to identify interspecies differences in ion signaling. We first identified orthologous human and yeast proteins in our datasets and found 2175 orthologs. As expected, these orthologs were enriched in proteins that are highly evolutionarily conserved, such as core metabolic pathway proteins, proteasome and ribosome^1^. Of these 2175 ortholog pairs, 1274 had no Ca^2+^-dependent thermal stability changes. For the proteins with Ca^2+^-dependent thermal stability shifts, 164 pairs of orthologs were identified as CRPs in both species, 203 were identified as CRPs only in human and 531 were identified as CRPs only in yeast (**Table S4**). These data suggest divergent cation engagement, and potentially regulation, of protein orthologs.

One major evolutionarily conserved pathway with a well-documented divergent Ca^2+^-regulation between human and yeast is the tricarboxylic acid (TCA) cycle^51,52^. In humans, the activity core TCA cycle enzymes oxoketoglutarate dehydrogenase (OGDH) and isocitrate dehydrogenase (IDH) is stimulated through a direct interaction with Ca^2+^, whereas *S. cerevisiae* lacks Ca^2+^ regulation of the TCA cycle^53,54^. We analyzed Ca^2+^-dependent thermal stability of yeast and human OGDH and IDH homologs to determine if our dataset captured this known divergent Ca^2+^ regulation of the TCA cycle. Ca^2+^ did not significantly engage the yeast Kgd1 (OGDH homolog) or yeast homologs of human IDHs, whereas human IDHs and OGDH were both significantly stabilized in the presence of Ca^2+^ ions (**Fig. 3A**). Moreover, this is in part explained by differential sequences within the conserved structures of human OGDH^55^ (PDB: 7wgr) and yeast Kgd1(AlphaFold)^56,57^. These structures aligned well overall (RMSD = 0.702Å; **Fig. 3B**), but the yeast Kgd1 lacks polar and acidic residues necessary for coordination of the Ca^2+^ ion corresponding to D154, D156, D158, and S160 in the Ca^2+^-binding pocket of human OGDH (**Fig. 3B**).

**Figure 3:**
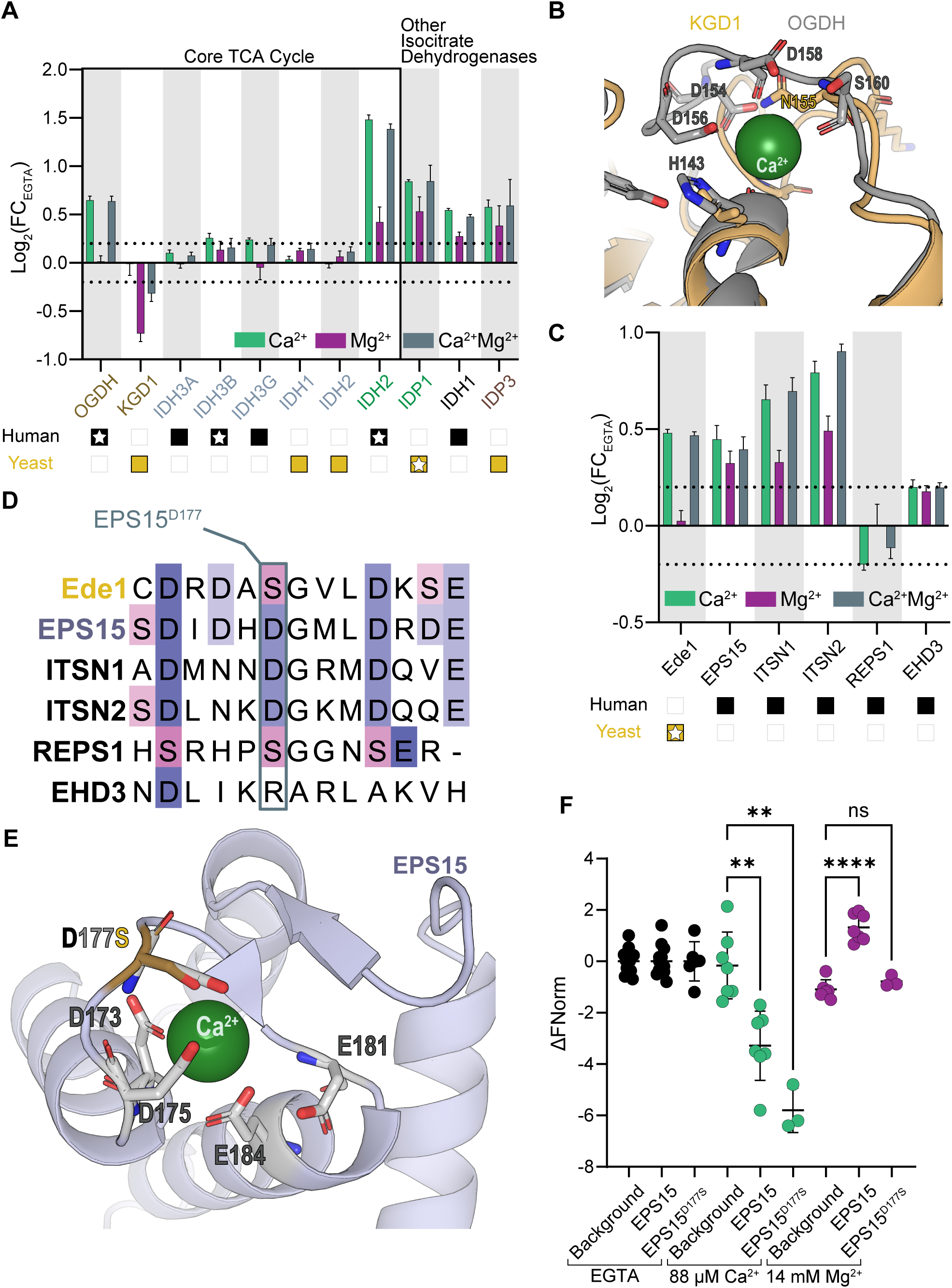
Ion engagement comparison of yeast and human proteomes captured evolutionary differences in Ca^2+^ signaling and ion coordination. **(A)** Ion-induced thermal stability changes of human and yeast OGDH and IDH homologs, relative to EGTA. Human TCA cycle dehydrogenases are CRPs, whereas yeast homologs do not engage Ca^2+^. Orthologs are indicated by the same color. Star denotes that the protein is a CRP in our dataset. Dotted lines show log_2_(FC) ± 0.2. Error bars: standard deviation of replicate measurements. **(B)** Overlay of human OGDH (gray) and its yeast homolog Kgd1 (yellow) structures at the Ca^2+^ coordination site. Yeast Kgd1 lacks residues that coordinate Ca^2+^ in human OGDH. **(C)** Ion-induced thermal stability changes of human and yeast EPS15 homologs, relative to EGTA. Yeast homolog Ede1 changes its abundance with Ca^2+^ addition only, human homologs EPS15, ITSN1, ITSN2 show thermal stability changes with Ca^2+^ or Mg^2+^. REPS1 and EHD3 do not show ion engagement. Dotted lines show log_2_(FC) ± 0.2. Error bars: standard deviation of replicate measurements. **(D)** Alignment of the second EF-hand domain of yeast and human EPS15 homologs. Yeast has a serine (S) where EPS15, ITSN1, ITSN2 have a conserved aspartic acid (D). REPS1 and EHD3 lack canonical EF-hand residues. **(E)** Overlay of EPS15^wt^ structure with a predicted EPS15^D177S^ structure. D177S mutation (stick representation, brown) is predicted to form a smaller coordination domain resulting in coordination of smaller Mg^2+^ ions. **(F)** Ca^2+^ or Mg^2+^ binding to purified EPS15^wt^ and EPS15^D1^^77^ proteins. Mammalian cells were transiently transfected with plasmids for expression and purification of EPS15-His proteins. Background refers to purification from cells that do not express tagged EPS15 proteins. Significance based on Student’s t-test: ns – not significant, ** - p<0.005, **** - p<0.0001.

Next, we investigated yeast-human homologous protein pairs with EF-hands in both homologs, but different thermal stability profiles with Ca^2+^ or Mg^2+^ addition. We identified yeast Ede1 and human EPS15 as one such pair. Ede1 thermal stability was Ca^2+^-dependent, but surprisingly, the human homologs EPS15, ITSN1, ITSN2, REPS1, and EHD3 either did not respond to cation addition or had significant responses to both Ca^2+^ or Mg^2+^. As a result, the human homologs were not considered as CRPs due to our hit criteria (**Figs. S1B, 3C**). To better understand the molecular basis of the ion-specific thermal stability differences between the yeast Ede1 and its human homologs, we aligned the second EF-hand domains of the homologs (**Fig. 3D**). Consistent with a lack of ion-induced thermal stability change, REPS1 and EHD3 do not have the canonical EF-hand domains required for ion coordination^58^. All of the acidic residues in the second EF-hand domain of Ede1 are shared with either EPS15 or ITSN1, with the exception of the Ede1 residue (S184) corresponding to D177 in EPS15 (**Fig. 3D**). Thus, we investigated whether mutation of EPS15 D177 to serine would increase EPS15 specificity for engaging Ca^2+^ (**Fig. 3E**). In this scenario, the larger aspartic acid group is predicted to form a binding cage to stabilize interactions with Mg^2+^, whereas the shorter serine side chain would only be able to coordinate and stabilize interactions with Ca^2+^ (ionic radius = 1.06 Å^59^), which is larger than Mg^2+^ (ionic radius = 0.81 Å^59^) (**Fig. 3E**). We mutated human EPS15 D177 to serine (D117S) to mimic the yeast EF-hand ion binding pocket and tested Ca^2+^ or Mg^2+^ binding to purified wild type (EPS15^wt^) and mutant (EPS^D117S^) proteins using microscale thermophoresis (MST) (**Figs. 3F, S3**). As expected, EPS15^wt^ and EPS15^D177S^ both bound Ca^2+^. However, the EPS15^D177S^mutant ablated Mg^2+^ binding, though Mg^2+^ binding was still observed for EPS15^wt^.

### Calcium regulation of DECR1 in murine mitochondria suggests that PUFA oxidation is a novel calcium-regulated mitochondrial metabolic pathway

Next, due to the importance of Ca^2+^ signaling in mitochondrial biology, we focused on analysis of the mouse data. We quantified 93.6% of the murine liver mitochondrial proteins annotated in MitoCarta^39^. Out of the 650 liver mitochondrial proteins identified, 182 passed our criteria with significant Ca^2+^-dependent altered thermal stability (**Table S3**). The submitochondrial localization and membrane association of these CRPs were representative of overall mitochondrial proteome (**Fig. S4A**). As noted above, we observed species specific cation binding effects in core TCA cycle proteins (**Fig. 3A**). In addition, an apparent Ca^2+^-regulation of DECR1, the rate limiting tetrameric enzyme of mitochondrial polyunsaturated fatty acid (PUFA) oxidation^26,27^ caught our attention (**Fig. 4A**). Surprisingly, the structurally conserved peroxisomal 2,4-dienoyl-CoA reductase DECR2 did not have Ca^2+^-dependent thermal stability changes (**Figs. S4C, S4D**). The mitochondrial fatty acid oxidation pathway has been shown to be regulated by Ca^2+^ signaling ^60^, but the mechanism of this regulation remains elusive. DECR1 is an attractive candidate involved in this regulation, as changes in DECR1 activity alter lipid metabolism^61,62^. To determine if the apparent Ca^2+^-dependent thermal stability shift of DECR1 is due to direct Ca^2+^-binding, we performed MST experiments with purified His-tagged human DECR1 protein **(Fig. S4B),** using Mg^2+^ as a control ion. Purified DECR1 bound Ca^2+^ but not Mg^2+^ (**Fig. 4B**). Unfortunately, no human structures of DECR1 with a bound Ca^2+^ ion exist. To identify potential Ca^2+^ binding pockets in DECR1, we aligned an empirical structure of DECR1 with a Ca^2+^ liganded predicted model of DECR1 from AlphaFill^17^. Based on a conserved binding pocket of a structurally similar bacterial dehydrogenase^63^, using a minimum sequence identity of 25%, AlphaFill predicted the site of Ca^2+^ binding within DECR1. This structural prediction suggests that Ca^2+^ aids in substrate recognition as the Ca^2+^ ion binds between the NADP+ and the substrate binding pockets^63^ (**Fig. 4C**).. In addition, alignment of this liganded model of DECR1 monomer and a tetrameric empirical structure of human DECR1 suggested that the Ca^+2^ could be coordinated adjacent to the NADP^+^ binding pocket, and that coordination was aided by E310 of a neighboring DECR1 molecule within the tetramer (**Fig. 4C**). Based on these models, we sought to determine whether DECR1 could bind Ca^2+^ at physiologically relevant concentrations and whether there was a substrate-dependent effect on Ca^2+^ affinity. Using MST, we measured the Ca^2+^ affinity of DECR1 in the presence or absence of 266 nM of NADPH and hexanoyl coenzyme A, a substrate mimetic (**Figs. 4D, S4E**)^26^. Without NADPH or substrate, DECR1 bound Ca^2+^ with a Kd of 12.0 ± 1.29 µM. In the presence of NADPH and hexanoyl coenzyme A, we observed an 8-fold reduction in DECR1’s Kd for Ca^2+^ (K_d_ of 1.48 ± 0.07 µM). Previous studies observed that the E310A mutation reduced the affinity of DECR1 for its substrate and NADPH by ∼2.8-fold each, and decreased enzyme activity by 50%^26^. Together with this work, our findings suggest a model wherein during a Ca^2+^ signaling event, DECR1 could use Ca^2+^ to regulate either substrate binding or substrate specificity and thereby enable Ca^2+^ signaling to fine tune metabolic outputs.

**Figure 4:**
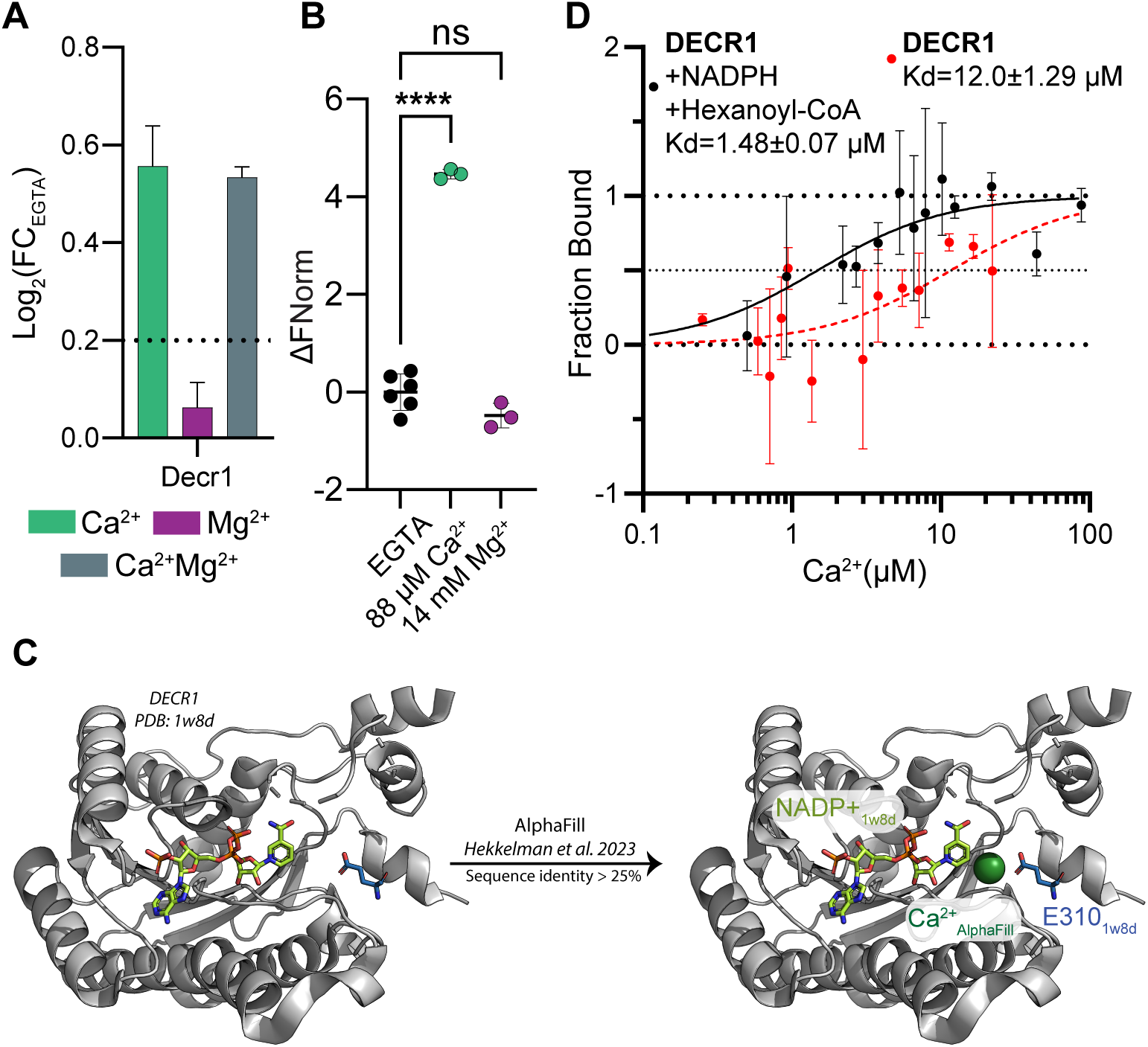
DECR1 is a novel Ca^2+^ binding protein. **(A)** Ion-induced thermal stability changes of mouse Decr1, relative to EGTA. Decr1 shows Ca^2+^-dependent stabilization. Dotted lines show log_2_(FC) ± 0.2. Error bars: standard deviation of replicate measurements. **(B)** Analysis of Ca^2+^ or Mg^2+^ binding to purified DECR1 using MST shows that DECR1 is a Ca^2+^ -binding protein. **** = p<0.0001. **(C)** Overlay of DECR1 structure with a Ca^2+^ liganded predicted model of DECR1 from AlphaFill. Placement of the Ca^2+^ ion (green) is based on a conserved binding pocket of a bacterial dehydrogenase that shows sequence homology to DECR1. The proximity of the Ca^2+^ ion to the NADP+ binding pocket and E310 of a neighboring DECR1 molecule within the tetramer suggests coordination. **(D)** Ca^2+^-binding curve for purified DECR1 using MST in the absence or presence of NADPH and hexanoyl-CoA (266 nM each). The Kd of DECR1 for Ca^2+^ is 1.48 ± 0.07 µM in the presence of its substrates, and 12.0 ± 1.29 µM in their absence.

### A resource to explore complex protein-cation interactions

We have shown the complexities of proteome-wide engagement by divalent cations in the context of divergent sequences and substrate affinities. The underlying data for this work is a rich source of new understanding of divalent cation regulation of diverse proteomes. From our initial analysis, we observed clear discrimination of proteome engagement with no cations, one cation, or two cations based on log_2_-normalized fold changes (**Fig. 1C**). We investigated this further by extracting the main factors driving discrimination of divalent cation engagement within the murine mitochondrial data. To do this we used the absolute values of log_2_ thermal stability changes because direct ligand binding can drive either an increase or decrease in thermal stability for a given protein; we then re-ran the PCA (**Fig. 5A**). PC1 separated the control samples from the cation treated samples, and PC2 separated samples based on the specificity of divalent cation binding (**Fig. 5A**) and extracted the loading for all mitochondrially-enriched, quantified proteins. As expected, using the PCA-based separation, we observed significant enrichment for Ca^2+^-binding proteins (**Fig. 5B, 5C**).

**Figure 5:**
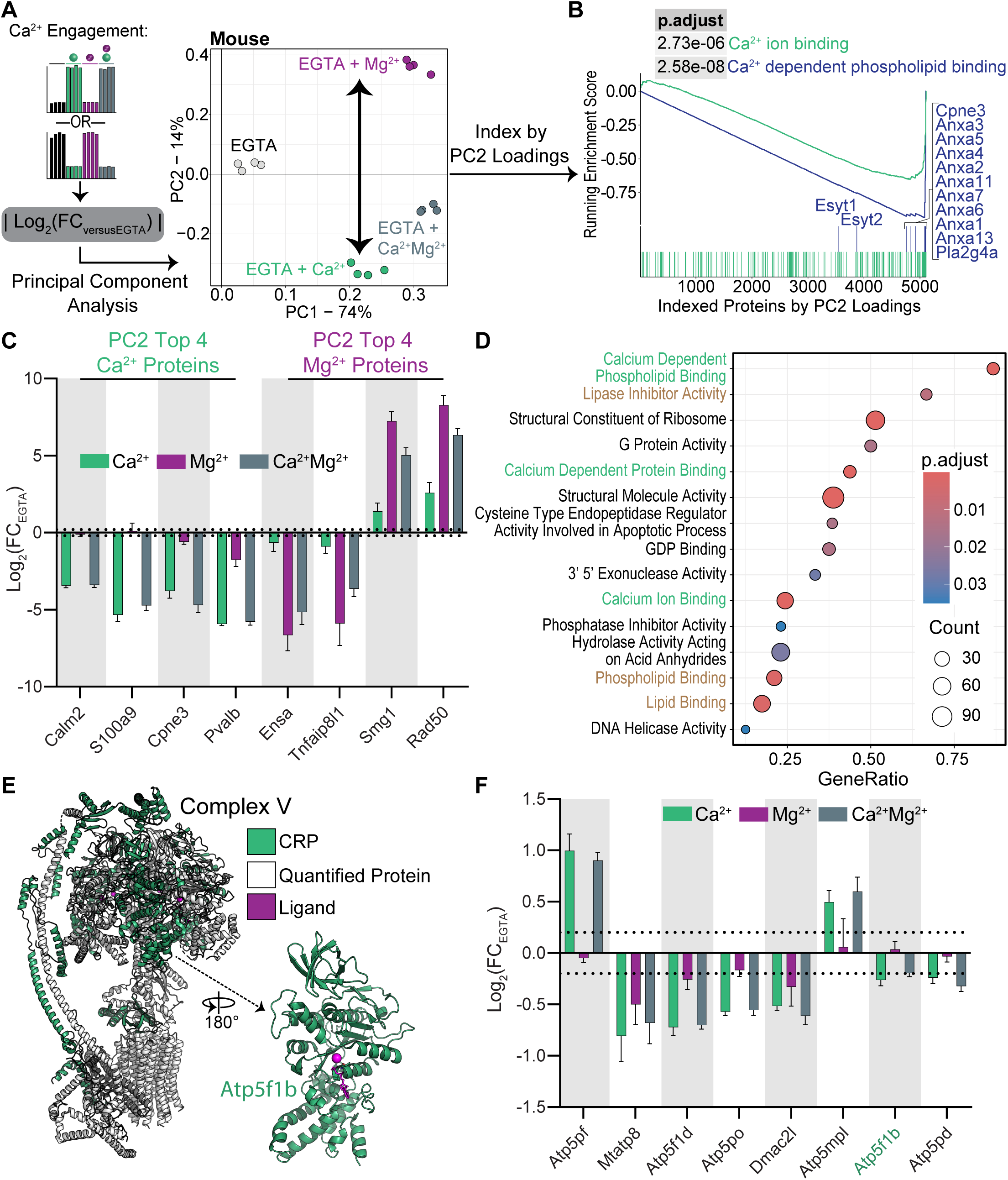
Exploration of calcium-specific engagement in murine mitochondria and oxidative phosphorylation complexes. **(A)** Workflow and output from PCA to assess Ca^2+^ or Mg^2+^ binding. |log_2_(FC)| was used based on the bidirectionality of thermal stability changes reflecting cation engagement. The resulting loadings were assessed for differentiation of the Ca^2+^ or Mg^2+^ conditions. PC2 separated proteins based on cation treatment and was used to index proteins in the subsequent analyses. **(B)** GSEA using PC2 loadings to index proteins showed strong enrichment for Ca^2+^ binding proteins at more negative loadings, consistent with separation of Ca^2+^ or Mg^2+^ binding specificity. **(C)** Cation-induced thermal stability changes of the top 4 proteins with the most negative and most positive PC2 loadings. Stability changes were consistent with specific engagement of either Ca^2+^ or Mg^2+^. Dotted lines show log_2_(FC) ± 0.2. Error bars: standard deviation of replicate measurements. **(D)** GSEA across PC2-loading space reported enrichment of multiple calcium binding protein gene sets (green text) as well as several lipid or membrane associated gene sets (brown text). **(E)** Mapping of mouse CRPs to the ATP synthase structure (PDB: 8h9v) revealed calcium engagement with the F1 domains, axle, and stator. ATP synthase ligands are shown in magenta. **(F)** Cation-induced thermal stability changes for ATP synthase complex proteins annotated as CRPs. CRPs were observed in all major domains of the complex. Dotted lines show log_2_(FC) ± 0.2. Error bars: standard deviation of replicate measurements.

Within the Ca^2+^-binding and, to a lesser degree, within the Ca^2+^-dependent phospholipid binding protein sets, we observed known Ca^2+^ binding proteins spread across the vector of PC2 loadings. In our initial analysis of the Ca^2+^-binding annexin proteins, we only investigated the 8 Ca^2+^-specific CRPs from the mouse mitochondrial data (**Fig. 2D**). While all of the 9 annexin proteins that we quantified in the murine mitochondria had large thermal stability changes with Ca^2+^ addition, Anxa13 had the largest magnitude thermal stability change with Mg^2+^ addition (**Fig. S5A**). From the PCA loadings discriminating between Mg^2+^ and Ca^2+^ engagement, we observed that Anxa13 thermal stability measurements were consistent with engagement of both divalent cations whereas Anxa3, for example, was highly specific for Ca^2+^ (**Fig. S5B**). Indeed, Anxa13 was thermally destabilized in the presence of Ca^2+^, and to a lesser degree with Mg^2+^ (**Fig. S5B**). Knowing that Anxa13 has been suggested to be the founder gene for the vertebrate annexins^64^, the lower Ca^2+^ specificity and failure to pass CRP hit calling for Anxa13 in our data may suggest that the other annexins became more specific for Ca^2+^.

Protein set enrichment of the PC-ordered murine mitochondrial data further revealed Ca^2+^-specific thermal stability profiles that were enriched for membrane associated complexes, lipase activity, and lipid binding (**Fig. 5D**). Based on this, we further investigated the Ca^2+^specific engagement of transmembrane proteins (**Fig. S5C**). PC-ordered separation of the transmembrane proteins identified cation-dependent thermal stability changes (**Table S1**). We further investigated the top two transmembrane proteins exhibiting thermal stability: Tmem165 and Tmem128 **(Fig S5C)**. Tmem165 was previously annotated as a putative divalent cation/proton antiporter human homologue of the yeast GDT1 and a known transmembrane Ca^2+^ transporter^65^. Tmem128 is not well characterized, though it contains an EF-hand-like sequence at its N-terminus (19-<u>E</u>V<u>E</u>L<u>D</u>C<u>E</u>D<u>E</u>AKP<u>E</u>-31). Additionally, in human cells, TMEM128 has been observed to interact with regulators of the endoplasmic reticulum such as reticulons RTN2 and RTN4 as well as REEP5 and REEP6 (**Fig. S5E**)^66,67^. In the murine mitochondria, we observed Reep6 as CRP demonstrating a role for Ca^2+^ regulation of Tmem128 and its ER-resident interacting partners (**Table S1**).

Finally, previous work has determined Ca^2+^ binding within the oxidative phosphorylation complexes^68^. While oxidative phosphorylation and the electron transport chain were enriched in PC2 space, these enrichments were not significant (**Fig. S6A**). We sought to further understand this lack of enrichment and modeled our observed CRPs on empirical structures for the oxidative phosphorylation complexes. Within the mitochondrial dataset, we observed a total of 26 oxidative phosphorylation complex members called CRP hits including 11 CI members, 2 CII assembly factors, 4 CIII members, 1 CIV protein, and 8 ATP synthase components (**Table S3, Fig. S6B)**. From these data we observed divergent effects with cation addition which explains the lack of separation in PC2 space. Within CI, Ndufa5 and Ndufa7 had Ca^2+^-specific thermal stability profiles, while Ndufb5 and Ndufa8 had profiles consistent with Mg^2+^-specific engagement (**Table S3, Fig. S6B**). We further mapped the observed CRPs to the protein structures for Complexes I and III-V (**Fig. 5E, S6D-F**). For example, we observed ATP5F1B, ATP5F1D within the F_1_ subcomplex of ATP synthase as CRPs, but not the other major member of the F_1_ subcomplex, ATP5F1A (**Fig. 5E,F**). These data are consistent with previous work that demonstrated direct binding of ATP5F1B to Ca^2+^ within the ATP/ADP binding pocket^69^, as well as conformational changes that take place along the ATP5F1D-ATPF1B axis in CV in the presence of Ca^2+^ during permeability transition pore (PTP) opening^70^. These data suggest that the observed Ca^2+^-dependent thermal stability effects happen at the level of specific proteins and their close interaction partners, rather than general destabilization of full complexes, and alludes to involvement of additional CV components in Ca^2+^-induced PTP opening. Moreover, visualization of CRPs in CI, CII and CIV revealed that proteins that engage other divalent cations (Mn^2+^, Cu^2+^, Zn^2+^, Mg^2+^), free FeS, and metal coordinating functional groups such as Heme were not identified as CRPs (**Fig. S6D-F)**, showing the specificity of thermal stability assay in identifying cognate metal-protein pairs.

## DISCUSSION

In this study we present a proteome-wide view of divalent cation engagement across three species. We generated high-throughput protein-metal interaction datasets detailing putative and known Ca^2+^-regulated proteins in human and yeast cells and extended these methods to characterize protein-metal engagement in whole respiring mitochondria. Across these datasets we quantified 2,884 proteins with significant thermal stability changes upon addition of Ca^2+^. Of these, 625 proteins were previously annotated as either being Ca^2+^-binding proteins or containing known Ca^2+^-binding domains. This high rate of recovery of Ca^2+^-binding proteins from an unbiased screen highlights the utility of thermal denaturation for the quantitative measurement of ligand binding interactions. In addition, we recovered proteins that are indirectly regulated by Ca^2+^ through their interaction with Ca^2+^-binding proteins. We conjecture that metal-dependent thermal denaturation can facilitate identification of signaling networks that utilize metal-regulated protein interactions. From these results, and the work of others ^20,21^, we believe that expanding these sets of non-specific protein-ligand interaction reference sets through studies such as thermal stability assays will enable comprehensive mapping of diverse and important classes of proteins as well as their key ligand regulators and signaling networks.

Interactions between Ca^2+^and Ca^2+^-binding domains such as the EF-hand domains and glutamate rich regions, have dissociation constants that can vary between nM to low mM values^71,72^. Because of this we chose to conduct our experiments at the upper end of this range. At low mM concentrations, we hypothesized that we would drive conformational and thermodynamic shifts for the broad range of potential metal binding proteins across all three species tested. To ensure that minimal residual Ca^2+^ or Mg^2+^ from cell lysates was present, we used 5 mM EGTA and were able to achieve depletion of cations to levels measured below their resting concentrations in cells.

Quantitative dissection of protein-ligand interactions has the potential to generate off-target engagement owing to changes in solution characteristics such as ionic strength. To control for changes in proteome thermal denaturation due to these factors, we performed quantitative comparisons between two divalent cations in each of our experiments. These analyses enabled the identification of Ca^2+^-specific protein stability changes as well as general effects caused by the addition of multiple divalent cations. Informatic comparison of putative Mg^2+^and Ca^2+^-binding proteins also helped us to determine site specific differences in protein domains between human and yeast cells that were responsible for altering general protein affinities for these metals.

Comparison of the yeast and human thermal proteome datasets led us to observe the difference in divalent cation specificity between Ede1 (yeast) and EPS15 (human) protein homologs. Interestingly, we were able to mutate a single site in EPS15 from an aspartic acid to a serine (D177S) that resulted in a drastic change in cation specificity, as mutant EPS15 could no longer coordinate Mg^2+^. EPS15 is a well-known substrate of the EGFR receptor tyrosine kinase and is involved in receptor mediated endocytosis to restrict EGFR activity ^73^. Previous work has shown that upon treatment with EGF, EPS15 localizes proximally to EGFR within 5 minutes of cell stimulation^74^. Thus, the combination of EGF stimulation-based EPS15 colocalization and cation selectivity hint at a potential means for human cells to integrate divalent cation sensing with receptor tyrosine kinase activity^58^ and even synaptic vesicle recycling^75^.

It has been known for several decades that Ca^2+^ regulates the TCA cycle in mammalian cells ^76^, in part by modulating the activities of pyruvate dehydrogenase phosphatase, isocitrate dehydrogenase, and alpha-ketoglutarate dehydrogenase ^77^.

We quantified components of all three of these complexes in the human dataset and two of them (OGDH1, IDH3B/G) were identified as CRPs. Conversely, we did not identify their yeast orthologs (KGD1, IDH1/2) as CRPs, consistent with lack of Ca^2+^ mediated activation of the TCA cycle in yeast. Comparative analysis of CRPs in different species has the potential to identify common elements and species-specific differences in Ca^2+^-signaling, such as the TCA cycle regulation. Here, we provide a rich dataset for such analysis to guide these efforts.

We identified site-specific modulation of divalent cation engagement in DECR1. These data highlight how large-scale analyses can be used to pinpoint individual sites. In addition, these data reinforce the importance of considering co-factor effects in thermal stability analyses. DECR1 was stabilized by Ca^2+^only when the thermal stability analysis was performed in situ (in murine mitochondria) and not in crude extracts (human analysis). Based on the MST binding experiments, we posit that this is due to the drastically reduced concentration of NADP/H (co-factor) and PUFAs in the human crude extracts. Indeed, predicted structures and Ca^2+^ binding sites within DECR1 suggest that Ca^2+^ is coordinated directly between NADP/H and the substrate binding pocket. Our follow up experiments using human DECR1 revealed µM affinity binding to Ca^2+^ that was increased 8-fold when substrate and cofactor were present. These data offer important clues for how the sensitivity for detection of Ca^2+^ binding proteins might be improved in the future and reinforce the need to consider the potential effects and requirements of secondary engagement when interpreting thermal stability experiments.

Analysis of thermal stability shifts of mitochondrial OXPHOS complexes showed calcium-dependent changes in individual proteins that tend to cluster or interact, rather than changes in the whole complex. We observed a similar pattern of individual proteins appearing as CRPs in the spliceosome. This observation evokes the possibility that through binding to select proteins, Ca^2+^ signaling can alter the function of OXPHOS complexes and the splicing machinery, and thermal profiling can be used to identify Ca^2+^ targets that are involved in such regulation. Consistent with this, we find two important components of mitochondrial CV for PTP function (ATP5F1B, ATP5F1D) as CRPs. An unexpected finding in our analysis was the strong enrichment of spliceosome components as CRPs. Although Ca^2+^-induced alternative splicing has been appreciated before ^49,50^, our data suggests a more general role for Ca^2+^ ions in RNA splicing.

Finally, our studies aimed to develop a resource to determine Ca^2+^-regulation across proteomes using unbiased proteomics methods. These data and our subsequent analyses hint at complex divalent cation regulation at the level of individual proteins, protein complexes, and organellar interfaces. Layered on top of these protein and localization considerations were factors such as the effects of additional ligands such as in the binding pockets of DECR1 and ATP5B. These data lay the groundwork for future studies to enhance our understanding of cation specificity and ligand dependencies within the context of proteome engagement of metal cations.

## MATERIALS AND METHODS

### Isolation of Mouse Liver Mitochondria

Protocols and reagents were adapted from Schweppe et al., 2017 and Frezza et al., 2007^30,31^. Livers were extracted from Bl6 adult mice and rinsed briefly in PBS twice before freezing in liquid nitrogen. Frozen livers were thawed in an isolation buffer (0.25 M sucrose, 20 mM HEPES sodium salt, 2 mM EGTA, pH 7.3) and minced into small pieces using scissors. Livers were then homogenized by a dounce homogenizer on ice and then centrifuged for 20 minutes at 800 x g at 4°C. The supernatants were collected and centrifuged for 20 minutes at 8,000 x g at 4°C. After the removal of supernatant, the pellet was resuspended in 30 mL isolation buffer. The two centrifugation steps were repeated twice. The protein concentrations of resuspended enriched mitochondria were quantified using a Bradford assay (BioRad). Enriched mitochondria were then aliquoted into 3.33 mg, centrifuged at 8,000 x g at 4°C, and frozen for future use.

### Mouse Mitochondria PISA Assay

Four conditions were prepared with the following biological replicates: 4x EGTA, 4x Ca, 4x Mg, and 4x Ca & Mg. Mitochondria were resuspended in PISA buffer (150 mM NaCl, 50 mM EPPS, 5 mM EGTA, pH 7.4) at a concentration of 2.66 mg/mL supplemented with a Roche protease inhibitor tablet and 1% digitonin (Sigma Aldrich). After 2 minutes, H_2_O was added to the EGTA condition, CaCl_2_ was added to the calcium condition to a final concentration of 10 mM, MgCl_2_ was added to the calcium condition to a final concentration of 5.6 mM, and CaCl_2_ and MgCl_2_ was added to the condition for final concentrations of 10 mM and 5 mM respectively. Two sets of 12 100 µL 2.66 ng/ µL enriched resuspended mitochondria aliquots in 200 µL PCR tubes were made per condition and heated on a temperature gradient of 42-64°C (2°C steps) for 5 minutes at each temperature using a Veriti 96-well thermal cycler (Applied Biosystems).

Samples were then left at 4°C for at least five minutes, the two sets were combined and transferred to 1.5 mL tubes, and then the samples were centrifuged for 20 minutes at 17,000 x g at 4°C. The supernatant was collected and combined for each condition, and soluble protein concentrations were then measured using a Bradford assay.

### Human and Yeast Whole Cell Lysate PISA Assays

For the human cell assay, HEK293T cells were cultured in RPMI supplemented with 10% FBS and 1% PenStrep, maintained in a 5% CO_2_ incubator at 37°C. 15cm plates of cells were harvested at 95% confluency with trypsin. The cells were then washed with PBS and cell pellets were collected and frozen in liquid nitrogen. For the yeast cell assay, *Saccharomyces cerevisiae* strain BY4742 (MATα his3Δ1 leu2Δ0 lys2Δ0 ura3Δ0) (Brachmann et al. 1998) was grown overnight in yeast peptone medium containing 2% glucose at 30°C. Overnight cultures were diluted to OD600 of 0.01 in the same medium and grown to an OD600 of 1 before cells were harvested by washing once in 1x yeast nitrogen base and freezing in liquid nitrogen.

Prior to the PISA assay, the respective cells were resuspended in lysis buffer (200mM EPPS, 150mM NaCl, 5mM EGTA, protease inhibitors, pH 7.2), then lysed using multiple freeze-thaw cycles using liquid nitrogen and thawing by vortexing to shear DNA. Lysate was cleared via centrifugation for 30 minutes at 21,130 x g. Supernatant was collected and stored on ice, then concentration was measured with a BCA assay. Lysate was then diluted to 2 mg/mL with more lysis buffer and divided into aliquots of 750µL each and kept on ice while treatment buffers were prepared.

Treatment buffers were prepared by adding stock solutions of 1M calcium chloride, 1M magnesium chloride, or both to the lysis buffer at twice the desired final concentration and held on ice. Each aliquot of protein lysate was then mixed 1:1 with respective treatment buffers and static incubated at room temperature for 15 minutes, then put back on ice. For each condition, 30µL of the respective lysate/treatment buffer mixture was aliquoted into each of forty PCR tubes in four sets of ten. Each set of tubes was held in an Eppendorf Mastercycler X50a thermocycler at 21°C for three minutes, then incubated for three minutes on a temperature gradient from 42 to 62°C. After the gradient, aliquots were equilibrated in the thermocycler at 21°C for three minutes. From each set of ten PCR tubes, 25µL of each aliquot was pooled for a total of four samples per condition and centrifuged for 1 hour at 21,130 x g. Finally, 25µL of each supernatant was immediately collected and frozen at -20°C overnight.

### Measurement of Ca^2+^ and Mg^2+^ Concentration in Lysates

Lysate samples for measuring free Ca^2+^ and Mg^2+^ concentrations were prepared as above but were aliquoted into a 96-well plate prior to the temperature treatment instead of proceeding with the PISA protocol. To measure free calcium concentrations of these lysates, either Oregon Green™ 488 BAPTA-6F, Hexapotassium Salt, cell impermeant (Invitrogen, O23990) or Fluo-4, Pentapotassium Salt, cell impermeant (Invitrogen, F14200) were used. The fluorescence values of these reads were compared to those read from calcium standards made with the Buffer Kit for Calibration of Fluorescent Ca² Indicators (Biotium, 59100). To measure free magnesium concentrations, the colorimetric Magnesium Assay Kit (Lsbio, LS-K220) was used. Experiments were performed following the manufacturer’s standard protocol with two biological replicates and three technical replicates using a Synergy H1 Microplate Reader (Agilent) to read fluorescence and absorbance.

### Sample Preparation and TMTpro Labeling

Each sample was reduced at a final concentration of 5mM DTT for 20 minutes at room temperature, then alkylated at a final concentration of 20mM iodoacetamide for one hour at room temperature, in the dark. IAA was quenched with 15mM DTT for 15 minutes at room temperature. Reduced and alkylated samples were then desalted with single-pot solid phase sample preparation (SP3) using Sera-Mag SpeedBeads. Bead-bound proteins were digested with a 1:100 ratio of LysC:protein overnight at room temperature, then a 1:100 ratio of Trypsin:protein for 6 hours at 37°C, each with gentle agitation. TMTpro labeling was done on-bead in 30% acetonitrile with a 2.5:1 ratio of TMT:peptide. After labeling at room temperature for 1 hour, a ratio check was done to check labeling efficiency before quenching the samples in 0.3% hydroxylamine.

### HPLC Fractionation and Mass Spectrometry Data Acquisition

Quenched samples were pooled and then desalted and resuspended in 5% acetonitrile, 10mM ammonium bicarbonate (pH 8.5). The sample was fractionated using basic-pH reverse-phase liquid chromatography, using a gradient from 5% acetonitrile/10mM ammonium bicarbonate to 90% acetonitrile/10mM ammonium bicarbonate. 96 fractions were collected over 75 minutes, which were then combined to make 24 fractions that were independently dried down and desalted. For the mouse mitochondria and human cell PISA, 12 alternating fractions were resuspended in 2% formic acid/5% acetonitrile and subsequently injected for analysis on an Orbitrap Eclipse Tribrid (Thermo Fisher Scientific). For the yeast cell PISA, all 24 fractions were analyzed. Peptides were separated using a 180-minute gradient on an in-house pulled C18 (Thermo Accucore, 2.6 Å, 150 μm) 30 cm column using an Easy nLC 1200 system (Thermo Fisher Scientific) running from 96% Buffer A (5% acetonitrile, 0.125% formic acid) and 4% buffer B (95% acetonitrile, 0.125% formic acid) to 30% buffer B. High-field asymmetric-waveform ion mobility spectroscopy (FAIMS) was enabled using compensation voltages of CV = -40/-60/-80V, “standard” resolution, and 4.6 L/min gas flow^78^. Spectral data was collected as follows.

MS1 scans were performed in the Orbitrap at 120,000 resolving power, 50 ms max injection time, and AGC target set to 100%. For each CV the top six precursors were selected for subsequent MS2 scans. Precursors selected from the MS1 scans were filtered based on intensity (min. intensity >5 × 10^3^), charge state (2 ≤ z ≤ 6), and detection of a monoisotopic mass (peptide monoisotopic precursor selection, MIPS). Precursor dynamic exclusion was set with a duration of 90 s, repeat count of 1, mass tolerance (10 ppm), and “exclude isotopes”. MS2 scans were collected using the linear ion trap, “rapid” scan rate, 50 ms max injection time, CID collision energy of 35% with 10 ms activation time, and 0.5 m/z isolation width. Finally, SPS-MS3 scans were collected at a resolving power of 50,000 with an HCD collision energy of 45%. Proteomics data is available through the ProteomeXchange/PRIDE with the identifier PXD048653.

### Mass Spectrometry Raw Data Processing

Raw files were converted to mzXML using Monocle^79^ and searched against the relevant annotated proteome from Uniprot (Human: October 2020; Mouse: March 2021; Yeast: October 2020). Common contaminant proteins and decoy protein sequences were appended to the beginning and end, respectively, of the Uniprot FASTA file. We used the Comet^80^ search algorithm to match peptides to spectra with the following parameters (unless noted all parameters were kept at default): 20ppm precursor tolerance, fragment_tolerance, variable_mods, static mods: TMTpro labels (304.207145) on peptide n-termini and lysine residues, alkylation of cysteine residues (57.0214637236), as static modifications, and methionine oxidation (15.9949146221) as a variable modification. PSMs were filtered to a 1% FDR at both the peptide and protein level using a linear discriminant analysis^81^. Protein-level FDR was filtered based on protein parsimony, and relative quantification of proteins was done using reporter ions.

### Proteomics Data Analysis

All data processing was performed in R (version 4.3.1) or Prism (10.1.2). Protein quantities were column normalized and used to calculate ratios relative to the mean for the control (first four) channels. Ratios were median-centered by column, then corrected using the COMBAT algorithm^82^. Unless otherwise noted, significance was assessed using a Welch’s t-test and these were corrected for multiple hypothesis testing using the Benjamini-Hochberg procedure^83^. Significant thermal stability changes were defined as a q-value of at most 0.05 and log_2_ fold change greater than 0.2 Proteins were determined to be CRPs based on Ca^2+^ engagement, a lack of Mg^2+^ engagement, and significant Ca^2+^ engagement even in the presence of Mg^2+^ (**Fig. S1B**). Gene ontology enrichment was performed using ShinyGO^35^. Gene set enrichment analyses were performed using clusterProfiler^84,86^ based on gene sets from MSigDB^87,88^.

### Cloning of DECR1 and EPS15, Mutagenesis

DECR1 cDNA (Genbank: BC105080.1) was obtained from DNASU Plasmid Repository. DECR1-6His was cloned into the mammalian expression vector pLJM1 (Addgene, 19319). Plasmid for expression of EPS15-GFP-His was obtained from Addgene (#170860). EPS15^D177S^ was generated using QuickChange Lightning Site-Directed Mutagenesis Kit (Agilent, 210518). All plasmids were sequence verified.

### Cell Culture, Protein Expression and Purification

HEK293T cells were grown in DMEM medium supplemented with 1× GlutaMAX (Gibco, 35050061) and 10% FBS (Avantor, 1300-500). The cells were tested for mycoplasma every 3 months using the Genlantis MycoScope PCR Detection Kit (VWR, 10497-508) and were confirmed to be free of mycoplasma contamination. The identity of the HEK293T cells was confirmed using short tandem repeat analysis. The HEK293T cell line has the following short tandem repeat profile: TH01 (7, 9.3); D21S11 (28, 29, 30.2); D5S818 (7, 8, 9); D13S317 (11, 12, 13, 14, 15); D7S820 (11); D16S539 (9, 13); CSF1PO (11, 12, 13); Amelogenin (X); vWA (16, 18, 19, 20); TPOX (11). This profile matches 100% to HEK293T cell line profile (CRL-3216; ATCC) if the Alternative Master’s algorithm is used, and 83% if the Tanabe algorithm is used.

HEK293T cells (∼5 million) were plated on 15 cm plates. The next day, cells were transfected with 5 µg of EPS15 or DECR1 expression plasmids using X-tremeGENE™ 9 DNA Transfection Reagent (Sigma-Aldrich, 6365787001) using 1:3 DNA:reagent ratio. 3 days after transfection, cells were lysed with 1% Triton-X-100 lysis buffer (50 mM HEPES KOH, 150 mM NaCl, 1% Triton-X-100, protease inhibitors, pH 7.2) and cleared via centrifugation for 10 minutes at 17,000 x g at 4°C. Lysates were then equilibrated and loaded into a tube with HisPur Ni-NTA resin (Thermo Scientific, 88221) per the manufacturer’s protocol. After 30-60 minute incubation at 4°C, the flow through was discarded and the resin was washed with modified HisPur wash buffers a total of six times: once with 0.5% Triton-X-100 added, once with 50 mM NaCl instead of 300 mM NaCl, once with 500 mM NaCl, and three with the wash buffer as listed by the manufacturer. For purifying DECR1-His, 150 mM imidazole was used. After washing, purified protein was eluted from the resin with elution buffer according to the HisPur purification protocol and then the purified protein was desalted using Pierce Zeba Desalting Spin Columns (Thermo Scientific, 89882) into MST buffer (20 mM HEPES, 150 mM NaCl, 7.5 µM EGTA, 0.05% Tween-20, pH 7.4) and frozen at -80°C.

### Western Blotting, SDS-PAGE, SYPRO Ruby Staining

SDS-PAGE and Western blotting as performed as described before^85^. The gels were stained with SYPRO Ruby Protein Gel Stain (Invitrogen, S12000) following manufacturer’s instructions and imaged with iBrightCL 1000 on fluorescent protein gel setting.

### Microscale Thermophoresis

Purified protein was labeled using the His-Tag labeling Kit RED-tris-NTA 2nd Generation (Nanotemper, MO-L018) following the manufacturer’s directions. Proteins were diluted to appropriate concentrations with MST buffer according to the affinity for the purified protein to the dye. Labeled proteins were mixed with either water, calcium (88 µM free Ca^2+^), or magnesium (14 mM free Mg^2+^) to check binding in the MST buffer. Binding affinity tests for DECR1-His were performed in the presence or absence of 266 nM of NADPH and hexanoyl coenzyme A each with free Ca^2+^ concentrations ranging from 11 nM-88µM. All MST experiments were performed using the Monolith under Nano-Red and 80% medium excitation intensity settings with an on-time of 1.5-2.5s. EPS15 experiments were performed using Regular Capillaries (Nanotemper, MO-K022), DECR1 experiments were performed using Premium Capillaries (Nanotemper, MO-K025) Analysis was performed using the MO Affinity software provided by the manufacturer.

## ACKNOWLEDGEMENTS

We thank the members of Zheng Lab at University of Washington for their help with MST measurements and instrument access as well as the members of the Schweppe and Villen labs for comments and technical advice for this work. This work was supported by the following funding sources: R35GM150919-01 (D.K.S), Andy Hill CARE Foundation (D.K.S.), Cancer Consortium New Investigator Award (D.K.S), R01GM138799-01 (D.M.S.), R01HL160825-01 (D.M.S.), Pew Charitable Trusts (Y.S.) and Royalty Research Fund of University of Washington (Y.S.).

## Supplemental Tables and Figures

**Supplementary Figure 1:**
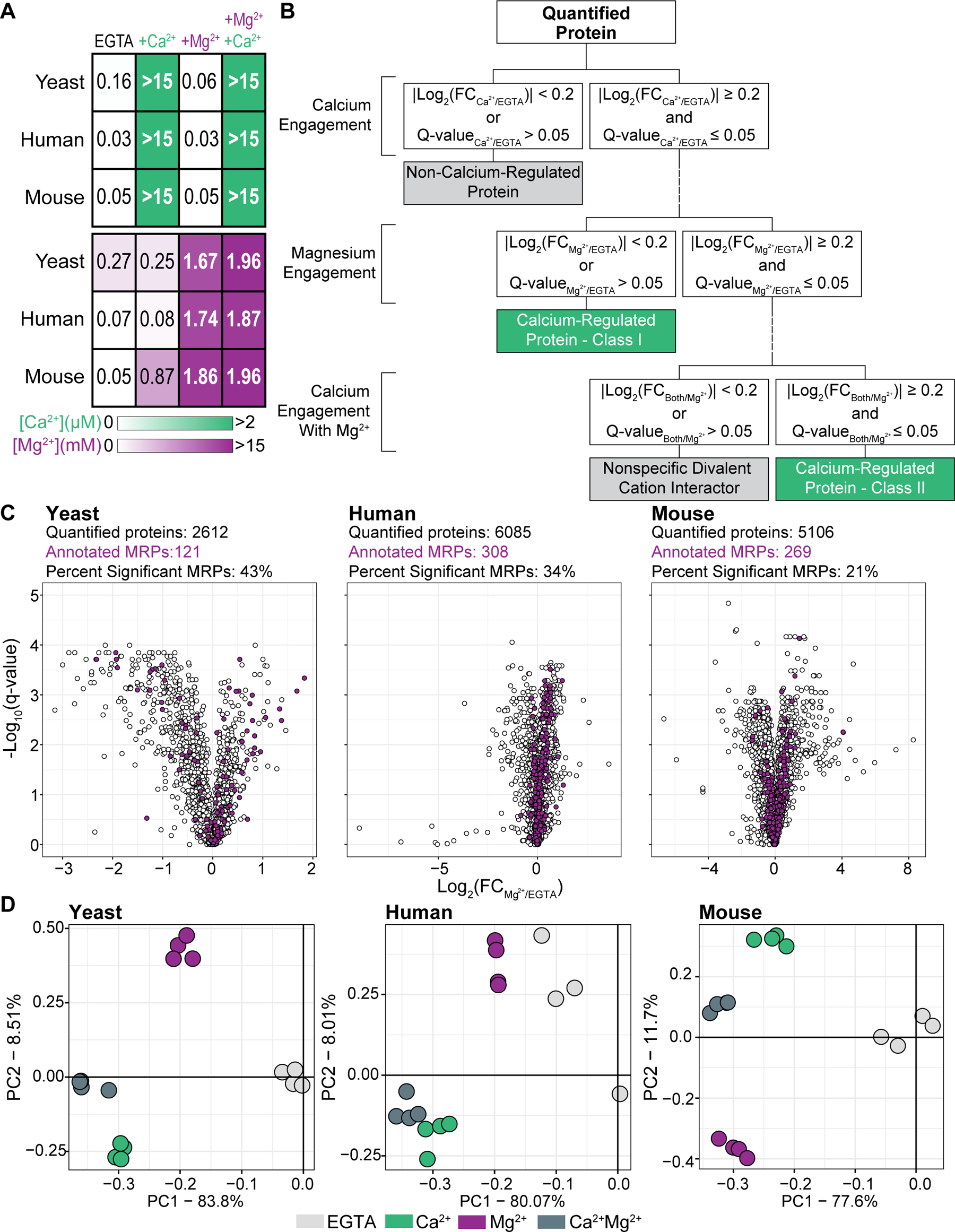
Details of PISA identification of Ca^2+^-regulated proteins. **(A)** Free Ca^2+^ and Mg^2+^ concentrations in lysates after EGTA or ion addition were measured using ion-specific fluorescent reporters. EGTA lowered concentrations of both ions to below physiological levels. Addition of ions increased their free concentration to physiological or above physiological levels. **(B)** Criteria for calling CRPs. Proteins were determined to be CRPs after satisfying a multilevel set of criteria based on significant thermal stability changes: (1) determination of Ca^2+^ engagement, (2) exclusion of non-specific ion engaging proteins (Mg^2+^-interaction), and (3) estimation of Ca^2+^ -specificity. Significant thermal stability changes satisfied the following two criteria: |log_2_(FC)| ≥ 0.2 and q-value ≤ 0.05. **(C)** Protein solubility changes in response to Mg^2+^ in three different proteomes. Volcano plots show protein abundance in Mg^2+^ relative to EGTA, and their q-values based on Welch’s t-test p-values corrected for multiple hypothesis testing. Proteins with |log_2_(FC)| ≥ 0.2, and q-value ≤ 0.05 were considered to show significant ion-dependent thermal solubility changes. Purple dots indicate proteins with previous Mg^2+^ annotation. **(D)** Principal component analysis of the yeast and human crude extract datasets as well as the mouse mitochondrial dataset. Points are colored based on treatment conditions with clear separation of different divalent cation conditions.

**Supplementary Figure 2:**
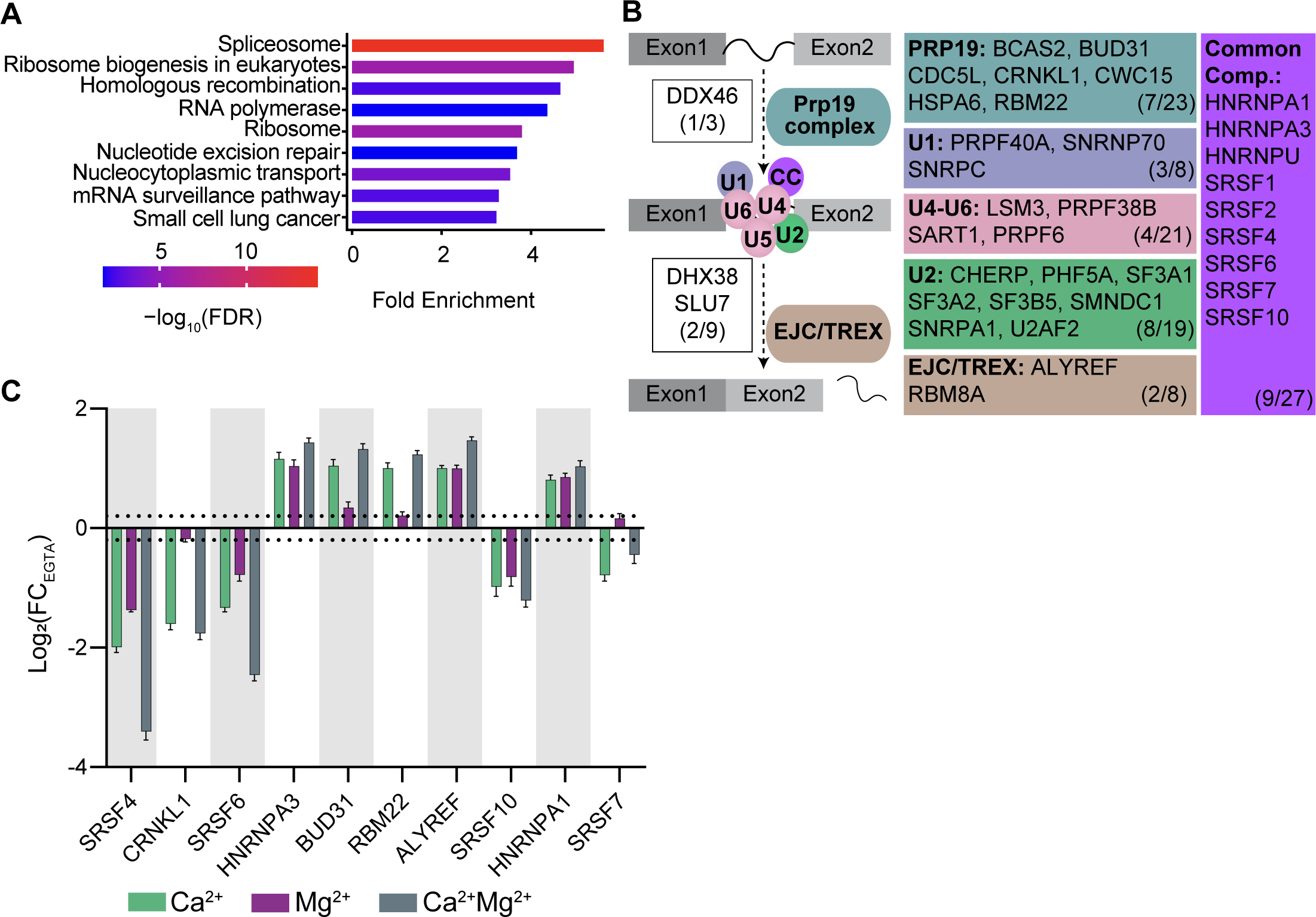
Spliceosome shows strong enrichment of Ca^2+^-regulated proteins. **(A)** GSEA human CRPs that do not have Ca^2+^ -related annotation. KEGG spliceosome pathway shows significant enrichment. **(B)** Schematic representation of a generalized splicing event (hsa:03040) and spliceosome proteins identified as CRPs in the human dataset. The protein components of each snRNP that were identified as CRPs are indicated in color matched boxes. White boxes indicate accessory non-complex proteins, and the purple box refers to common components (CC) present throughout splicing. The number of CRPs or the number of component proteins quantified are indicated in color matched boxes. **(C)** Ion-induced thermal stability changes of 10 example human spliceosome CRPs, relative to EGTA. Dotted lines show log_2_(FC) ± 0.2. Error bars: standard deviation of replicate measurements.

**Supplementary Figure 3:**
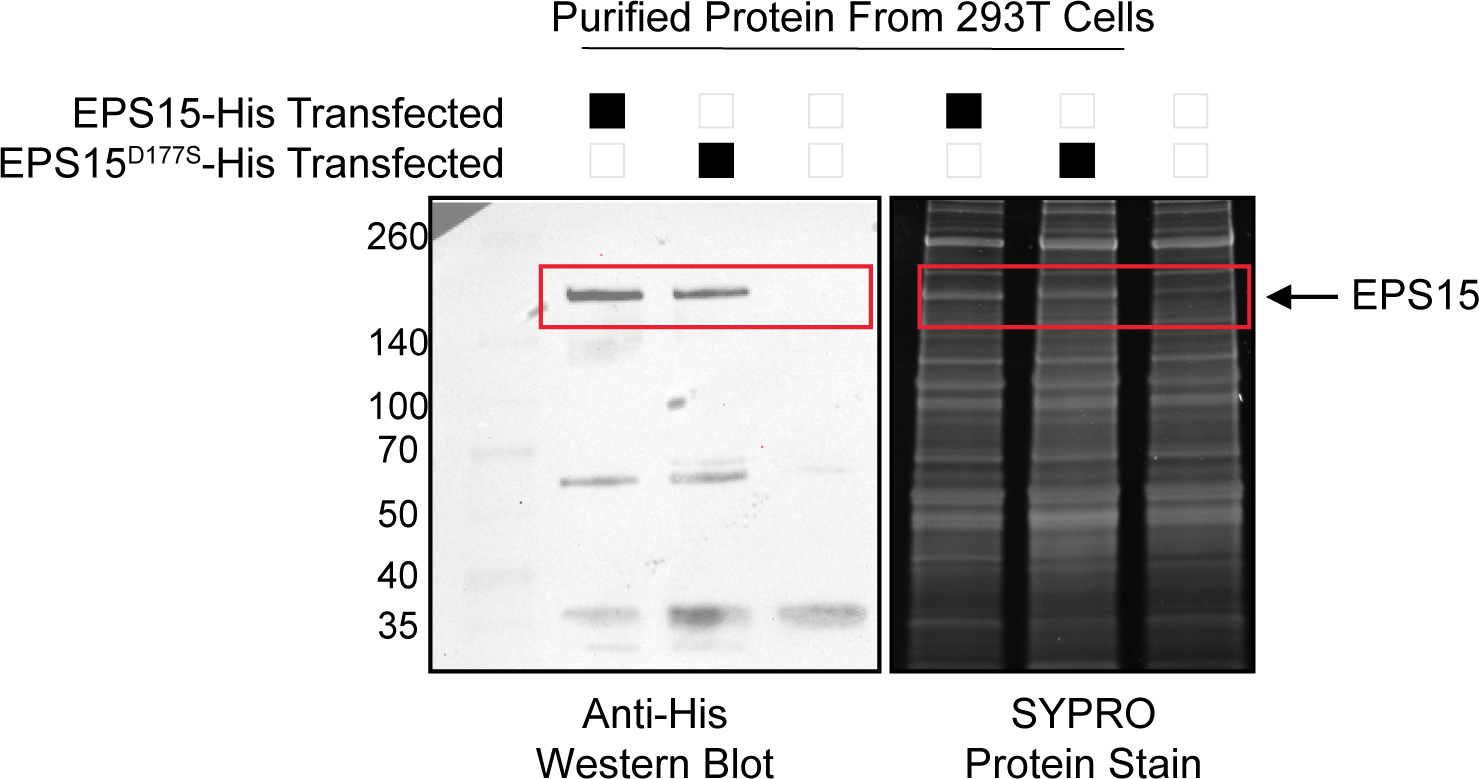
Purification of EPS15^WT^ and EPS15^D177S^ from HEK293T cells after transient transfection. Western blot (left) and SYPRO Ruby protein stain (right) analysis of EPS15 purifications. Cells that do not express His-tagged EPS15 were used as controls.

**Supplementary Figure 4:**
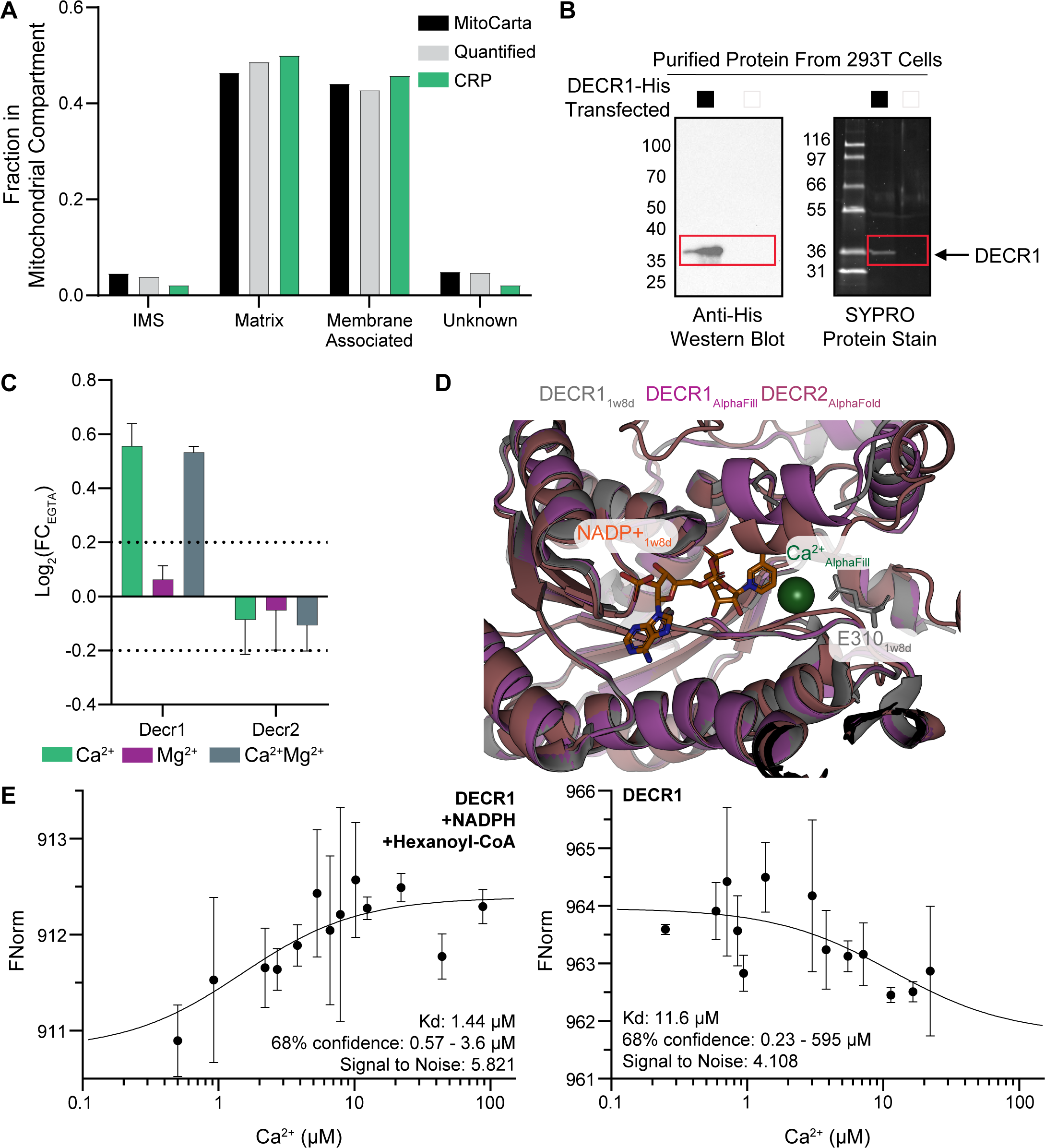
Analysis of mitochondrial CRPs. **(A)** Submitochondrial localization of MitoCarta proteins, proteins quantified in mouse experiments, and mouse CRPs are shown. Mitochondrial CRPs are identified in all submitochondrial compartments. **(B)** Western blot (left) and SYPRO Ruby protein stain (right) analysis of DECR1 purification. Cells that do not express His-tagged DECR1 were used as controls. **(C)** Ion-induced thermal stability changes of mouse Decr1 and Decr2, relative to EGTA. Decr1 shows a significant Ca^2+^-dependent thermal stability change whereas Decr2 does not. Dotted lines show log_2_(FC) ± 0.2. Error bars: standard deviation of replicate measurements. **(D)** Overlay of DECR1 structure with a Ca^2+^ liganded predicted model of DECR1 from AlphaFill and DECR2 structure from AlphaFill. DECR2 has structural similarity to DECR1, but the binding pocket is structurally different with no E310 from a neighboring DECR2 molecule to coordinate calcium. **(E)** Ca^2+^ binding curves of DECR1 in the presence (left) and absence (right) of NADPH and hexanoyl-CoA using MST.

**Figure S5:**
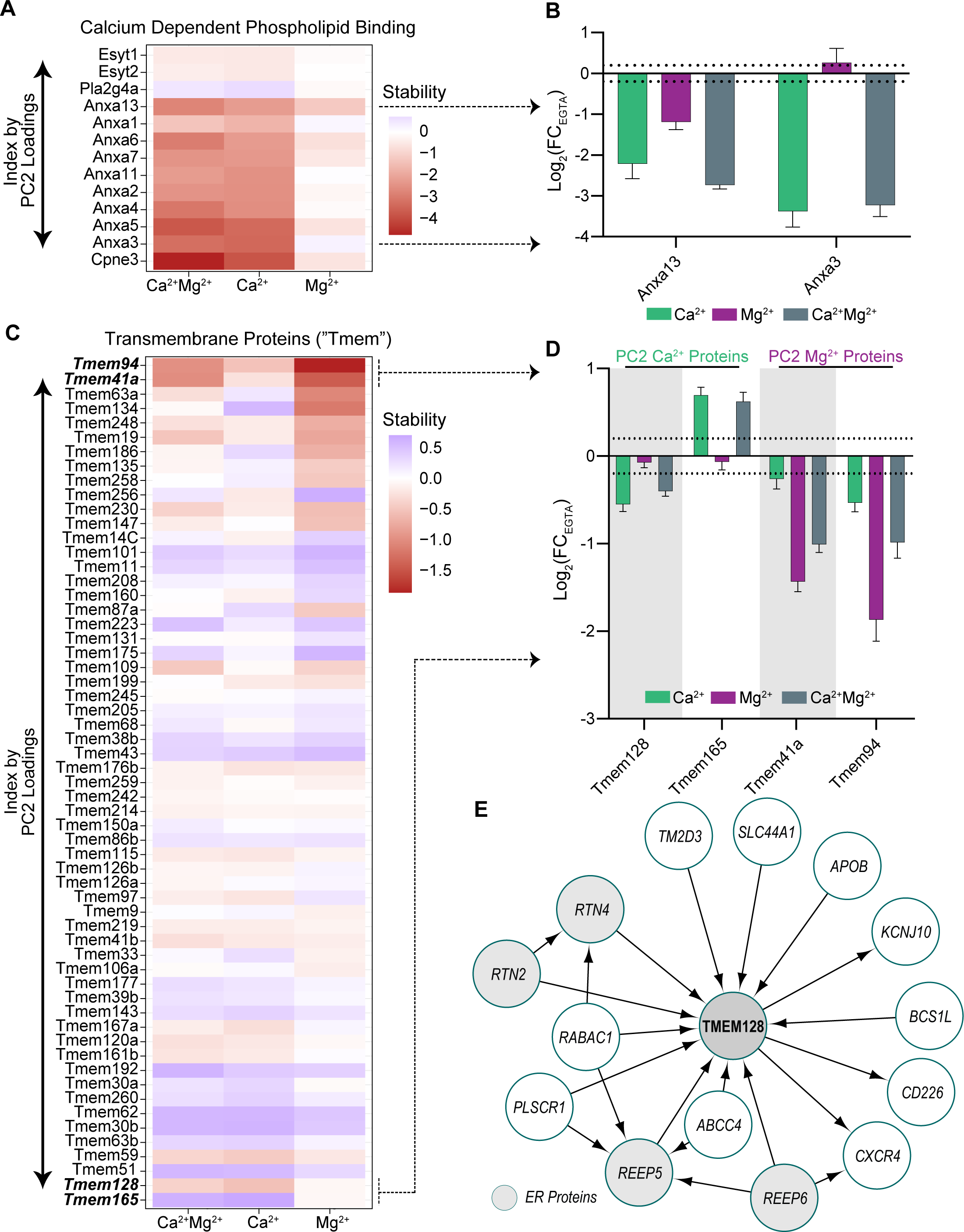
Cation engagement of calcium dependent phospholipid binding proteins and transmembrane proteins. **(A)** Based on the GSEA shown in Figs. 5B and 5D, plotting the relative thermal stability compared to control samples for the GO Molecular Function set of ‘Calcium dependent phospholipid binding’. Proteins are ordered by PC2 loadings from the PCA in Figure 5. **(B)** Highlighting the relative thermal stability compared to control samples for Anxa3 and Anxa13. Anxa13 had the highest level of Mg^2+^ thermal stability alteration, and therefore the lowest Ca^2+^ specificity. Dotted lines show log_2_(FC) ± 0.2. **(C)** From the GSEA in Figure 5D, proteins along the PC2 loadings were enriched for lipid binding and membrane associated functions. Based on these findings we investigated transmembrane proteins (“Tmem”) along PC2 and observed divergent engagement with cations for these proteins. **(D)** From S5C, the top two ‘calcium specific’ proteins we observed were Tmem128 and Tmem165. The top ‘magnesium specific’ proteins we observed were Tmem41a and Tmem94. Dotted lines show log_2_(FC) ± 0.2. **(E)** Tmem128 protein interaction network established from the BioPlex interactome. Tmem128 interacts with at least four known endoplasmic reticulum (ER) proteins: RTN2, RTN4, REEP5, REEP6 (highlighted in grey).

**Figure S6:**
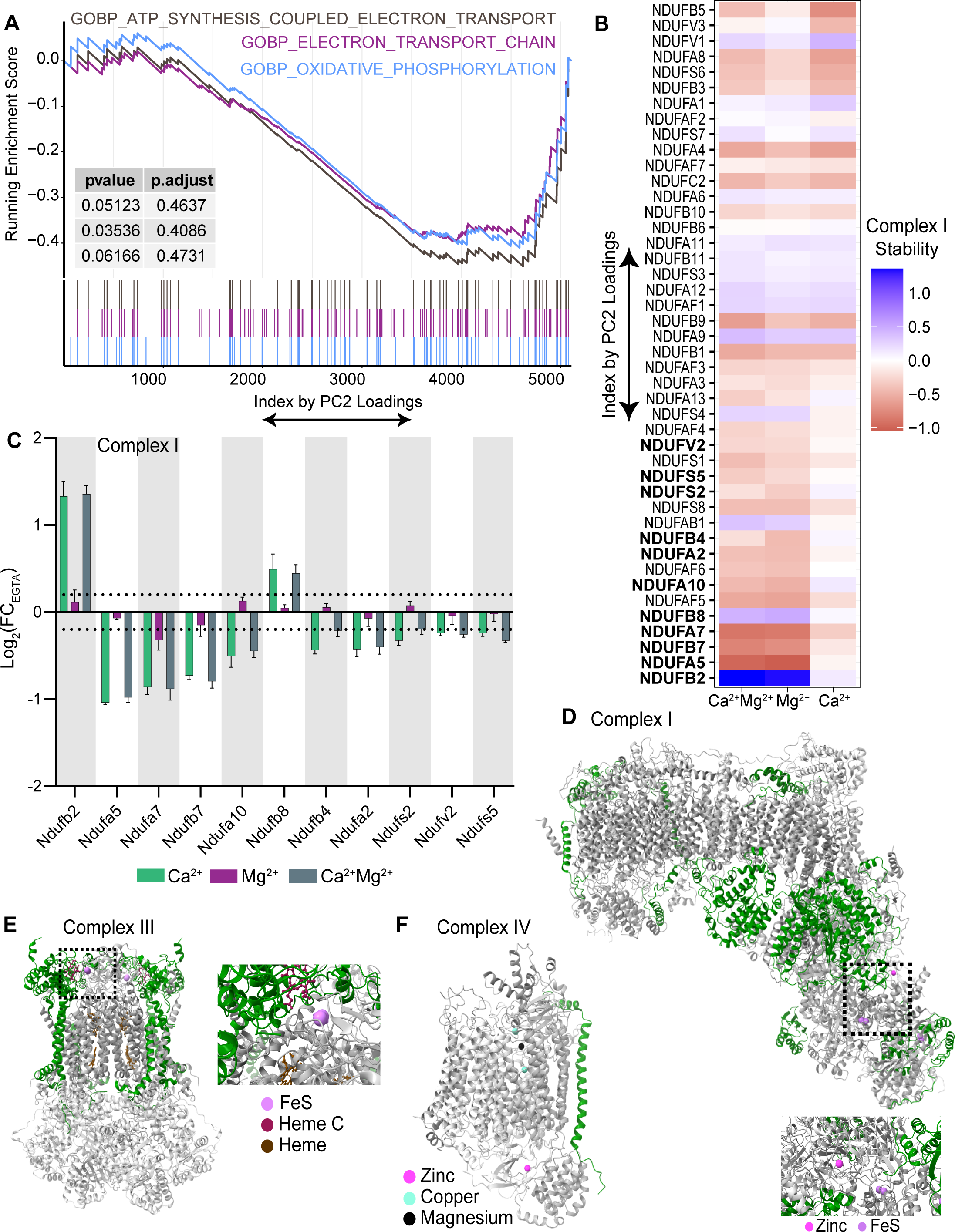
Cation engagement within the oxidative phosphorylation complexes. **(A)** GSEA results for terms related to oxidative phosphorylation were not significantly enriched based on the PC2 loadings from murine mitochondrial proteins. **(B)** Relative thermal stability compared to control samples for proteins in Complex I. Proteins were ordered based on their PC2 loadings from PCA in Figure 5. Complex I stability colored based on log_2_(FC) compared to EGTA condition. **(C)** Relative stability changes of the 11 Complex I proteins that were called as CRPs in murine mitochondria. Dotted lines show log_2_(FC) ± 0.2. Error bars: standard deviation of replicate measurements. **(D)** Overlay of the 11 Complex I CRPs onto the structure of Complex I (PBD: 5LNK). Observed CRPs (highlighted in green) were not in direct contact with known cations or ligands. Iron sulfur (FeS) highlighted in lavender, zinc highlighted in magenta. **(E)** Overlay of the 4 Complex III CRPs onto the structure of Complex III (PBD:5XTE). Observed CRPs (highlighted in green) were not in direct contact with known cations or ligands. Iron sulfur (FeS) highlighted in lavender, heme C highlighted in red, heme highlighted in gray. **(F)** Overlay of the 1 Complex IV CRPs onto the structure of Complex IV (PBD:5Z62). Magnesium highlighted in gray, zinc highlighted in magenta, copper highlighted in blue.

